# Hidden introns and repeated gains shape the heavily interrupted tRNA genes of Asgard archaea

**DOI:** 10.64898/2026.10.13.758758

**Authors:** Satwato Goswami, Akshoy Kumar Ghosh, Krishnendu Sinha

## Abstract

Transfer RNA (tRNA) genes are usually compact, yet in some Asgard archaea—the closest known prokaryotic relatives of eukaryotes—most are interrupted by introns. Studying this has been hampered by circularity: archaeal introns are catalogued by software that finds them partly through the splicing motif later used to explain where they persist. We built a motif-free census that detects intervening sequences by comparing each tRNA gene with compact relatives, and applied it to 90 Asgard genomes and 18 reference archaea. The standard annotator, tRNAscan-SE, had missed 30% of intervening sequences in the three focal families, mostly in the D-arm, but almost none in reference archaea; six independent checks supported the census. After correction, focal tRNA genes carry 2.2 intervening sequences on average, and in 30 genomes chosen before analysis the two focal families ranked first and second of 21 (exact *P* = 1*/*210). A reconstruction based on the census rather than on annotated introns placed 129 high-confidence gains at 65 loci on 37 branches of the species tree, 71 of them terminal; the two intron-rich classes are not sister lineages and carry mostly different intron positions, consistent with independent accumulation. Intervening sequences sat at canonical bulge–helix–bulge splicing motifs six to ten times more often than the average of three matched null models, including those the annotator missed and the reconstructed gains. Tests of copying between tRNA genes, mobile elements and reverse transcription found no general route of intron arrival. Asgard tRNA introns were thus gained repeatedly and sit preferentially at sites predicted to allow splicing; how they arrive remains open.

**Significance statement:** Every cell needs transfer RNAs to make proteins. In some Asgard archaea, the microbes most closely related to the ancestors of animals, plants and fungi, almost every tRNA gene is broken up by introns, which must be cut out before a tRNA can work. We show that the standard software for finding these introns misses nearly a third of the candidate intron sequences, and we provide a simple, motif-free way to find them. With a corrected catalogue, the evidence shows that introns were inserted many separate times, apparently independently in two lineages, and that they sit preferentially where the RNA can fold into the shape that the cell’s splicing enzyme is predicted to recognise. The study also shows how a detection tool can bias evolutionary conclusions about the very feature it detects, and how to test for that bias directly.

## 1 Introduction

Transfer RNAs (tRNAs) translate the genetic code. Each is a molecule of about 76 nucleotides that folds into a cloverleaf and then into a compact L-shape, and nearly every part of that fold has a job: the acceptor stem carries the amino acid, the anticodon reads the messenger RNA, and the D- and T-arms hold the structure together. It is therefore surprising that tRNA genes can be interrupted by introns—stretches of sequence that are copied into the precursor RNA and must be removed before the tRNA can work (Fig. 1a). In archaea and eukaryotes the classic tRNA intron sits at one “canonical” position in the anticodon loop, between nucleotides 37 and 38 (Abelson et al., 1998; Marck and Grosjean, 2003; Yoshihisa, 2014). Archaea also carry “noncanonical” introns at many other positions, and some archaeal tRNA genes contain two or more (Marck and Grosjean, 2003; Sugahara et al., 2007; Randau and Söll, 2008).

**Figure 1.**
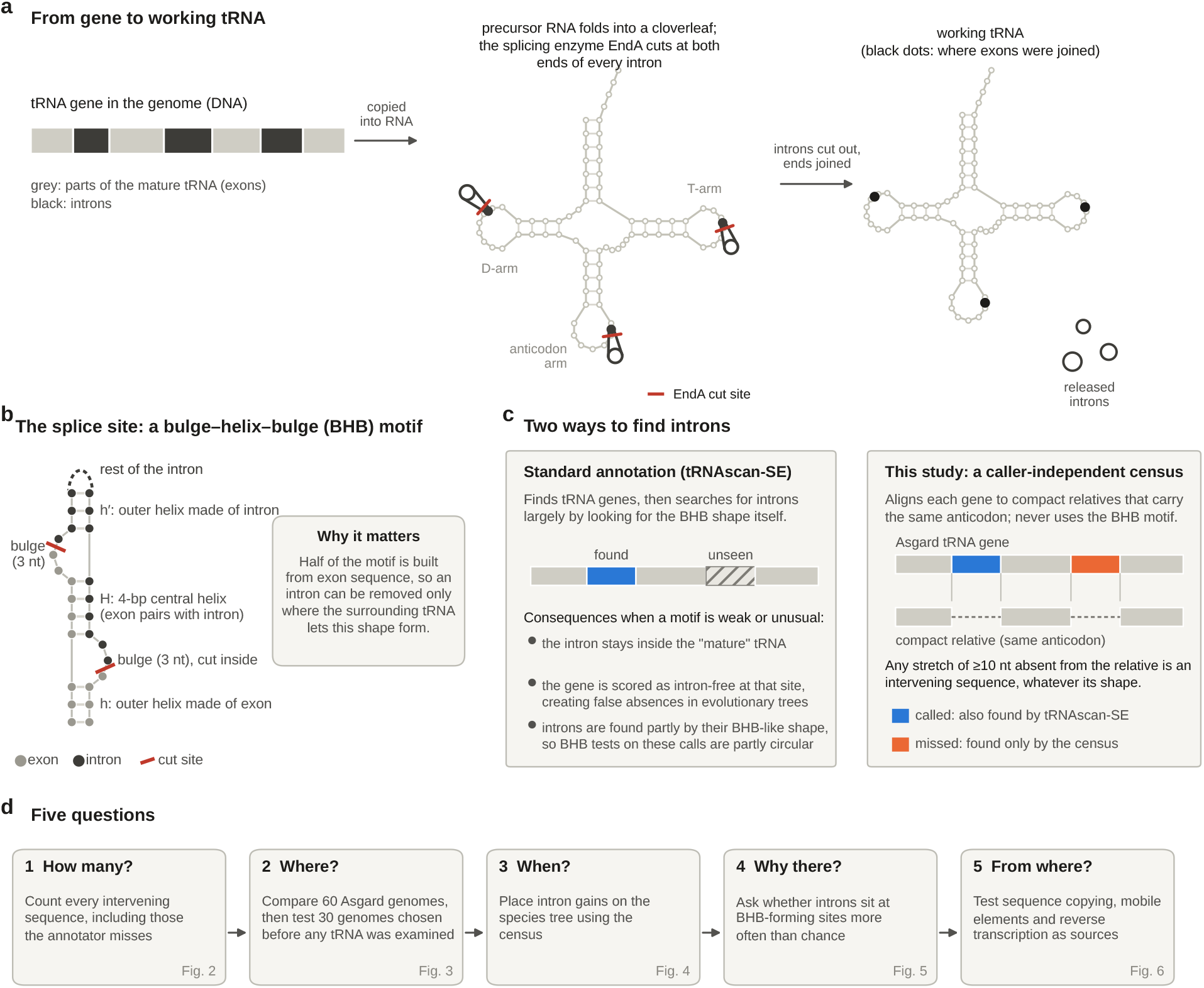
tRNA introns, the motif that allows their removal, and the problem of finding them. **a**, A tRNA gene interrupted by introns (black) is copied into a precursor RNA that folds into a cloverleaf. The splicing endonuclease EndA cuts at both ends of each intron (red bars), the exon pieces are joined, and a working tRNA is released. **b**, The bulge–helix–bulge (BHB) motif at an intron boundary. Two three-nucleotide bulges sit on opposite strands of a four-base-pair central helix (H), flanked by outer helices made of exon (h) or intron (h^*′*^) sequence; EndA cuts inside each bulge. Because part of the motif is exon, an intron can be removed only where the surrounding tRNA allows the motif to form. **c**, The standard annotator finds noncanonical introns largely by searching for the BHB shape, so introns with a weak or unusual motif can be left inside the “mature” tRNA, the gene is scored as intron-free at that position, and any BHB test on the calls is partly circular. The census used here compares each gene with compact relatives of the same anticodon and flags every extra stretch of *≥*10 nucleotides regardless of shape. Wherever annotation status is shown, blue means called by tRNAscan-SE and orange means missed by tRNAscan-SE and found only by the census; other colours are defined in each legend. **d**, The five questions addressed and the figures that answer them.

Archaeal tRNA introns are removed by the splicing endonuclease EndA, which recognises a local RNA shape called the bulge–helix–bulge (BHB) motif: two three-nucleotide bulges on opposite strands separated by a four-base-pair helix, with one cut in each bulge (Fig. 1b) (Marck and Grosjean, 2003; Tocchini-Valentini et al., 2005; Hirata et al., 2012). Because half of the motif is built from exon sequence, an intron can be removed only where the surrounding tRNA allows the shape to form. This makes tRNA introns a direct window on how RNA structure constrains genome evolution. In the archaeal order Thermoproteales, for example, related intron sequences recur at different positions and in different tRNA genes, which has been interpreted as large-scale intron transposition constrained by the BHB requirement (Sugahara et al., 2008; Fujishima et al., 2010; Tocchini-Valentini et al., 2011).

Asgard archaea are the prokaryotes most closely related to eukaryotes (Spang et al., 2015; Zaremba-Niedzwiedzka et al., 2017; Imachi et al., 2020; Eme et al., 2023; Zhang et al., 2025). Three complete genomes from wetland soil revealed an extreme tRNA architecture: 92–95% of tRNA genes contained an intron, more than half contained several, and many introns were noncanonical (Valentin-Alvarado et al., 2024). Notably, the authors found additional introns by inspecting secondary structures with R2DT (RNA 2D Templates; Sweeney et al., 2021) that the standard annotator had not reported (Valentin-Alvarado et al., 2024). These observations raise five questions. How many introns do these genes really carry? How widespread is the phenotype? Did it arise in a single ancestral burst or through many separate gains? Why do the introns persist where they do? And how does intron DNA enter a tRNA gene in the first place?

Two methodological problems stand in the way (Fig. 1c). The first is circularity. The standard annotator, tRNAscan-SE, identifies noncanonical archaeal introns largely by searching for BHB-like structures (Chan et al., 2021). Any test of whether introns sit at BHB motifs that uses the same calls is therefore partly testing the software. Introns that the software misses are not only uncounted; they are silently scored as *absences*, and such false absences can create spurious evolutionary “gains”. The second problem is non-independence. Most Asgard genomes are metagenome-assembled drafts, near-identical genomes are common, and neighbouring intron calls can share a single evolutionary origin, so raw counts overstate the amount of independent evidence.

Here we address both problems directly. We built a census that detects intervening sequences without using the BHB motif, by comparing each tRNA gene with compact relatives that lack the insertions, and checked it in six independent ways. We used it to correct the intron catalogue, to test whether the phenotype recurs in genomes chosen before any tRNA was examined, to reconstruct intron gains without relying on annotation, and to test the BHB association on sequences that the annotator never saw as introns. Finally, we asked whether any single route of intron arrival explains the pattern (Fig. 1d).

## 2 Results

### 2.1 Standard annotation misses nearly a third of Asgard tRNA intervening sequences

Many “mature” Asgard tRNAs are too long. A class I tRNA is normally 70–78 nucleotides long without its 3^*′*^ CCA end. Yet after the introns annotated by tRNAscan-SE were removed, 48% of class I tRNA genes in the three focal families—Freyrarchaeaceae, Jordiarchaeaceae and Atabeyarchaeaceae—were still at least 88 nucleotides long, compared with 16% in other Asgard families and fewer than 1% in 18 reference archaea (Supplementary Fig. S1a). In tRNAscan-SE’s own secondary-structure output these extra stretches appear as unpaired insertions, most often in an enlarged D-loop.

To count such sequences without relying on the BHB motif, we aligned every tRNA gene, with all of its sequence retained, to compact tRNAs of the same anticodon drawn from all 108 genomes in the study (Fig. 2a). Every stretch of at least 10 nucleotides present in the gene but absent from its best-matching compact relative was recorded as an intervening sequence, and then labelled *called* or *missed* according to whether tRNAscan-SE had annotated it. The census never uses the splicing motif.

**Figure 2.**
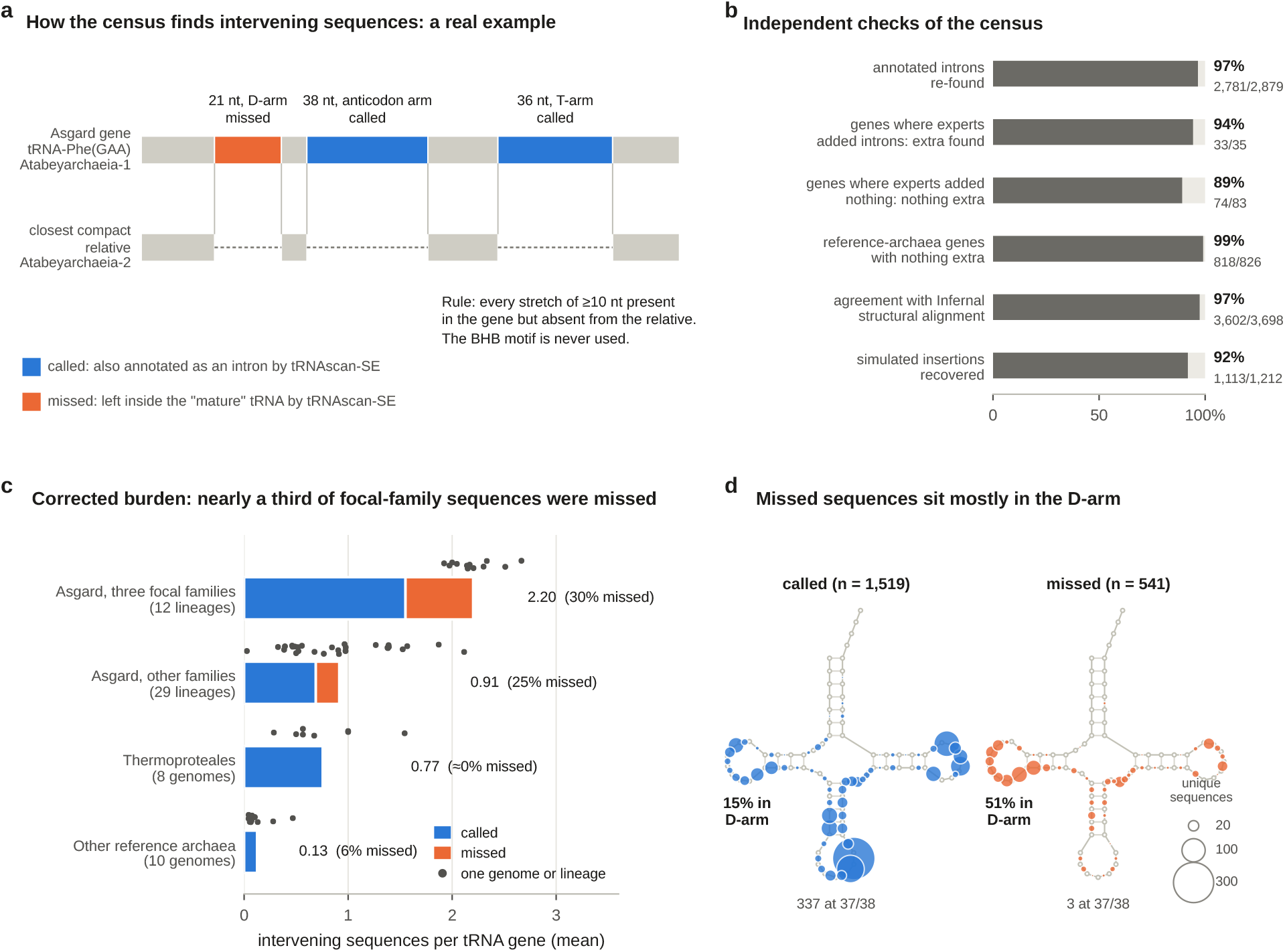
A caller-independent census shows that standard annotation misses nearly a third of Asgard tRNA intervening sequences. **a**, A real example. The tRNA-Phe(GAA) gene of the complete Atabeyarchaeia-1 genome is aligned to the closest compact tRNA-Phe(GAA), from the complete Atabeyarchaeia-2 genome. Three stretches are present only in the first gene: tRNAscan-SE annotated two of them (blue) but left a 21-nucleotide D-arm insertion inside the “mature” tRNA (orange). **b**, Independent checks of the census: recovery of annotated introns; agreement with expert curation of the three complete genomes, both where curators added introns and where they added none; absence of extra sequence in reference archaea; gene-by-gene agreement with an Infernal covariance-model alignment; and recovery of simulated insertions. The sixth check, a model-free length comparison, is in Supplementary Fig. S1a. **c**, Intervening sequences per tRNA gene, split into called (blue) and missed (orange). Bars are group means; dots are individual genomes (reference archaea) or lineages (Asgard; near-identical genomes merged, patristic distance *<*0.005 substitutions per site). **d**, Where intervening sequences sit on the tRNA cloverleaf (discovery panel; one point per distinct sequence, area proportional to the number of sequences at that position). Called introns concentrate at the canonical anticodon-loop position and in the T-arm; missed sequences concentrate in the D-arm. Class II variable-arm insertions are omitted from the drawing.

Six independent checks support the census (Fig. 2b; Supplementary Fig. S1). It re-found 2,781 of the 2,879 introns that tRNAscan-SE had called in the discovery panel (97%). It found almost nothing extra in reference archaea: 8 intervening sequences in 826 tRNA genes, including only 4 beside the 277 called introns of eight intron-rich Thermoproteales genomes. It agreed with expert curation of the three complete Asgard genomes: where the authors of the genome study had added introns to the automated calls (Valentin-Alvarado et al., 2024), the census found extra sequence in 33 of 35 genes, and where they added nothing it found nothing in 74 of 83. It agreed gene by gene with an entirely different method, an Infernal covariance-model alignment of the “mature” sequences (Nawrocki and Eddy, 2013): the two methods agreed on whether a gene carried a missed insertion for 97% of 3,698 genes, and the amounts of inserted sequence were highly correlated (Pearson *r* = 0.95). Finally, in a simulation in which real intervening sequences were inserted at random into compact tRNA genes, the census recovered 92% of the insertions, placed the boundary within three nucleotides in 92% of those recovered, and showed no blind spot in the D-arm (94% recovery there; Supplementary Fig. S1c,d).

Applied to the discovery panel, the census found 1,235 intervening sequences that tRNAscan-SE had missed, in 947 of 2,456 tRNA genes; the independent panel contained a further 423 in 341 of 1,242 genes.

Correcting for these omissions makes the phenotype more extreme (Fig. 2c). After merging near-identical genomes into 12 distinct lineages, tRNA genes in the three focal families carry on average 2.20 intervening sequences each, compared with 1.55 annotated introns; 30% of their insertions had been missed. In a typical focal lineage, 97% of tRNA genes carry at least one intervening sequence and 71% carry two or more (annotation alone: 87% and 53%), with up to five in a single gene. For comparison, the most intron-rich reference genome, *Pyrobaculum calidifontis*, has 1.54 introns per tRNA gene, all detected by tRNAscan-SE. Other Asgard families carry fewer intervening sequences (0.91 per gene on average), of which 25% had been missed (missed fractions are ratios of group means). Requiring insertions of at least 13 nucleotides gives the same picture (Supplementary Fig. S2a).

Missed and called sequences were alike in length (median 22 and 23 nucleotides; Supplementary Fig. S2b) but not in location (Fig. 2d). Half of the 541 distinct missed sequence–host combinations (51%) lay in the D-arm, compared with 15% of the 1,519 distinct called ones, whereas the canonical anticodon-loop position was almost never missed (3 distinct sequences). The annotator therefore systematically overlooks D-arm insertions—precisely the noncanonical positions that make Asgard tRNAs unusual. This reflects intron architecture rather than assembly quality: across 90 Asgard genomes the fraction missed rose with the number of annotated introns per tRNA gene (Spearman *ρ* = 0.47, *P* = 3 *×* 10^*−*6^) but was unrelated to contig number or N50 (|*ρ*| *<* 0.02) and only weakly to completeness (*ρ* = 0.13, *P* = 0.21; Supplementary Fig. S2c). Because related genomes are not independent, we repeated these correlations after merging near-identical genomes (71 units): the association with annotated burden remained (*ρ* = 0.37, *P* = 0.001), and none with assembly metrics appeared (|*ρ*| *≤* 0.10, *P ≥* 0.41). Per-gene census results and every intervening sequence are listed in Supplementary Table S3 and Supplementary Table S4.

### 2.2 Heavy interruption is concentrated in two classes and recurs in genomes chosen before analysis

Across the discovery panel, high burden is concentrated in the three focal families and is consistent within each of them (Fig. 3a). These families belong to two classes of the Genome Taxonomy Database (GTDB), Jordiarchaeia (Freyrarchaeaceae and Jordiarchaeaceae) and Atabeyarchaeia, which are not each other’s closest relatives on the species tree. The census also exposes a burden that annotation had hidden elsewhere: the single Borrarchaeaceae lineage carries 2.12 intervening sequences per tRNA gene, close to the focal families, but half of them (51%) had been missed. Sifarchaeaceae (1.87) and DXJG01 (1.57) carry elevated burdens as well, whereas several Heimdallarchaeia and Njordarchaeia lineages carry fewer than one intervening sequence per gene.

**Figure 3.**
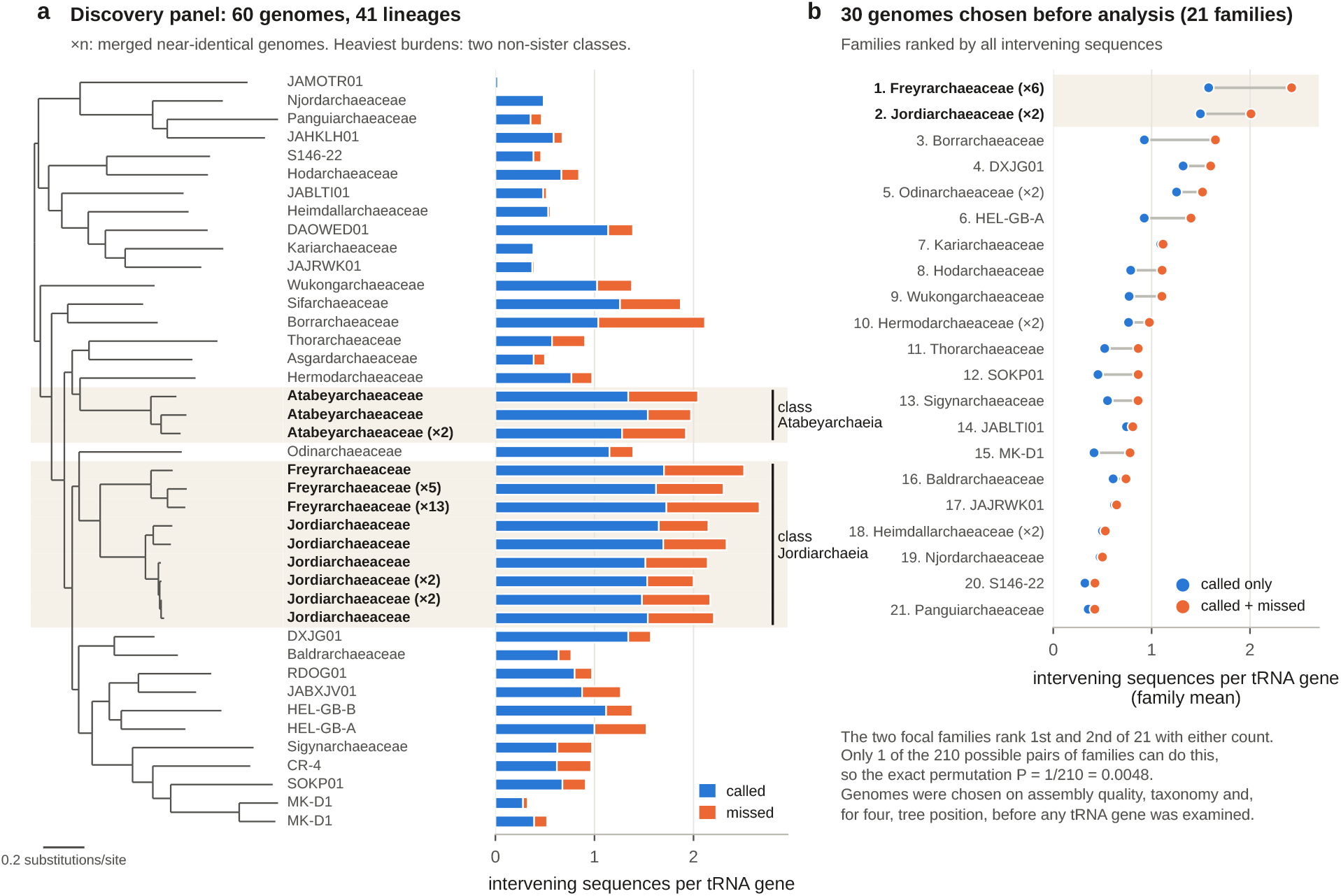
Heavy tRNA interruption is a property of two classes and recurs in genomes chosen before any tRNA was examined. **a**, Species tree of the 60-genome discovery panel (GTDB 53-marker maximum-likelihood tree, midpoint-rooted), with near-identical genomes merged into 41 lineages (*×n*: number of merged genomes). Bars show intervening sequences per tRNA gene: called by tRNAscan-SE (blue) and found only by the census (orange). Shading marks the three focal families, which fall in two classes (brackets) that are not sister groups. **b**, The 30-genome independent panel, summarised by family and ranked by all intervening sequences (*×n*: number of genomes). Blue dots, annotated introns only; orange dots, annotated plus missed. The two focal families rank first and second of 21 under both measures; the exact permutation probability of this outcome is 1*/*210.

To test whether this pattern was an accident of which genomes we had studied, we analysed 30 additional assemblies selected on assembly quality, taxonomy, representative status and, for four non-focal genomes, proximity to the focal families on the species tree, with the list fixed before any tRNA was examined (Supplementary Table S1). The panel contained six Freyrarchaeaceae and two Jordiarchaeaceae genomes and 22 genomes from 19 other families. Whether we counted annotated introns or all intervening sequences, the two focal families ranked first and second of the 21 families (Fig. 3b). Freyrarchaeaceae and Jordiarchaeaceae averaged 2.42 and 2.01 intervening sequences per tRNA gene, against a median of 0.86 (maximum 1.65, Borrarchaeaceae) for the other families. There are exactly 210 ways to choose two families out of 21, and only one of them—the true focal pair—places both at the top, so the exact permutation probability is *P* = 1*/*210 = 0.0048, the smallest value this design can produce (Supplementary Table S6). Within the focal independent genomes, 98% of tRNA genes carry at least one intervening sequence and 74% carry two or more. Because all eight focal genomes are draft assemblies, these data establish that the phenotype recurs, not how common it is in every member of either family (Supplementary Note 4).

### 2.3 Introns were gained many times, apparently independently in the two intron-rich classes

To reconstruct history without depending on annotated introns, we rebuilt the evolutionary character matrix from the census. We first matched tRNA genes across genomes into corresponding loci using sequence, structure and conserved neighbouring protein-coding genes. Within each of the 172 multi-genome loci that carried an insertion, we mapped every intervening sequence onto a common coordinate system and grouped insertions whose mapped boundaries lie within three nucleotides into a single intron position, giving 541 intron positions (characters). A genome was scored as having an intron at a position, lacking it (its copy of the locus was recovered and had no insertion nearby), or unknown (the locus was not recovered). No tRNAscan-SE intron call entered these states.

An intron counts as a high-confidence gain when it is present in a well-supported group of genomes on the species tree (ultrafast bootstrap *≥*80), absent from no recovered genome inside that group, and absent from at least one good-quality relative outside it. This rule identified 129 gains (hereafter census gains) at 65 loci on 37 different branches (Fig. 4a; Supplementary Table S8). Most are recent: 71 lie on terminal branches.

**Figure 4.**
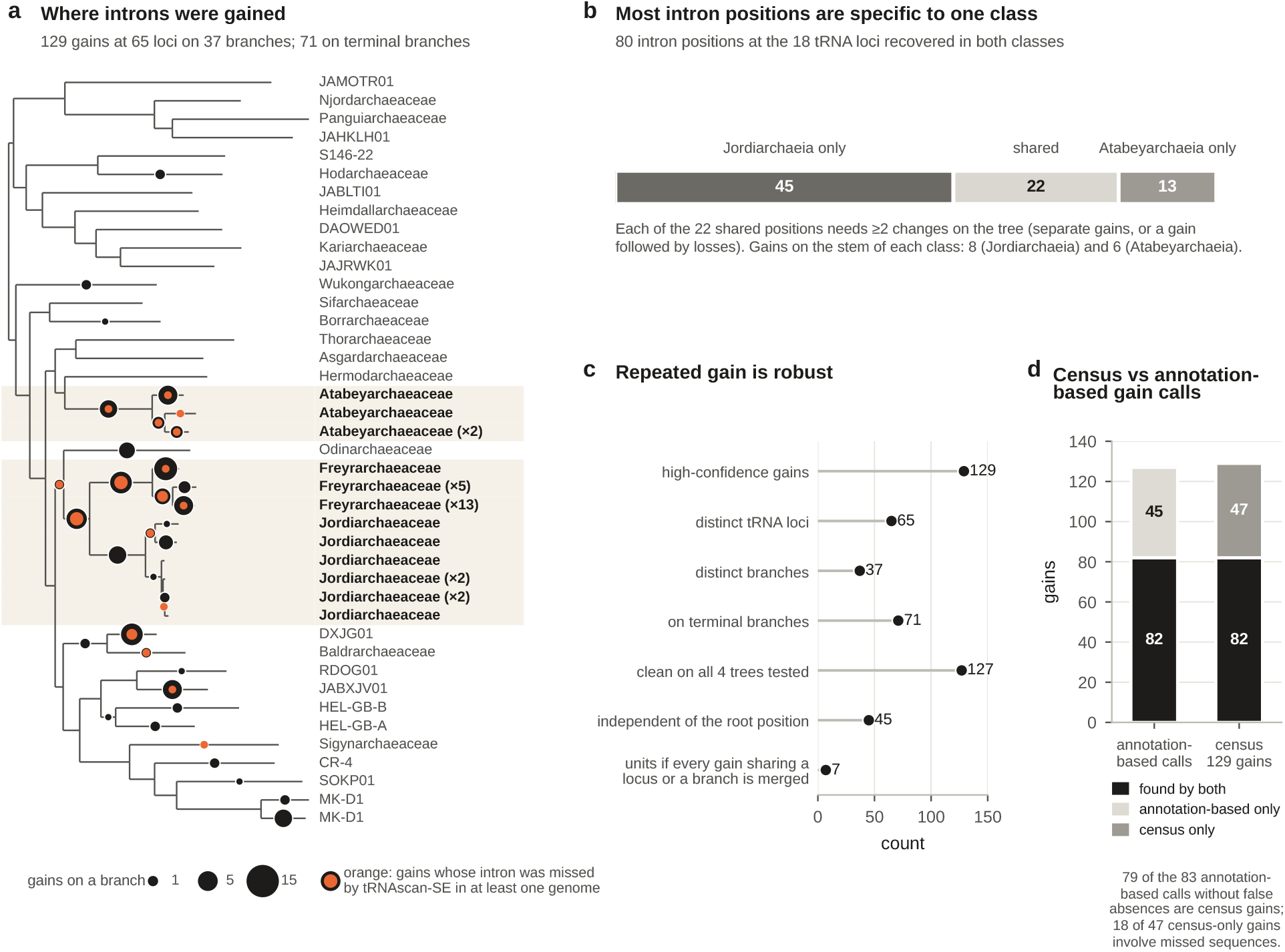
Intron gains reconstructed from census intron states. **a**, High-confidence gains on the species tree (lineages as in Fig. 3a; one of the 37 gain branches lies inside a merged lineage and is drawn on its tip). Circle area is proportional to the number of gains on that branch; the orange inner circle marks the share of those gains whose intron was missed by tRNAscan-SE in at least one genome. **b**, Intron positions at the 18 tRNA loci recovered in both intron-rich classes. Most positions occur in only one class; each of the 22 shared positions requires at least two changes on the tree. **c**, Robustness of the reconstruction: distinct loci and branches carrying gains, gains on terminal branches, gains that remain clean on all four trees tested (project tree, project tree with branches of support *<*95 collapsed, GTDB releases 226 and 232; only the internal gains on the GTDB trees are informative, see text), gains independent of the root position, and the number of units that remain if every gain sharing a locus or a branch is merged. **d**, Comparison with gains reconstructed from tRNAscan-SE calls alone. Of 127 annotation-based calls, 82 are census gains; most of the remainder rest on false absences (Supplementary Fig. S7). Of 129 census gains, 47 are not among the annotation-based calls.

Two observations argue against a single ancestral burst. First, the two intron-rich classes are not sister groups: their most recent common ancestor is also the ancestor of 14 other families in our panel (26–27 on the GTDB reference trees), most of which carry fewer introns (Fig. 3a); a burst in that ancestor would require losses in many intervening lineages. Second, at the 18 loci recovered in both classes, 58 of 80 intron positions occur in only one class and 22 are shared (Fig. 4b). Every shared position requires at least two changes on the tree—separate gains, or an ancient gain followed by losses in other lineages—so none is explained by a single gain in the common ancestor alone. Gains on the stem branch of each class (8 in Jordiarchaeia and 6 in Atabeyarchaeia), on many internal branches and on terminal branches show that introns accumulated throughout the history of each class. More broadly, 104 of the 541 intron positions changed state at least twice on the tree (Supplementary Fig. S8c), so the same positions were often gained or lost repeatedly in different lineages.

These conclusions do not depend on the tree used. Terminal gains do not depend on tree topology, and every gain clade has ultrafast-bootstrap support of at least 95, so collapsing weakly supported branches of the project tree leaves all 129 intact. The informative test of topology is therefore on the 58 internal gains: on the GTDB release 226 and 232 reference trees, which differ in deep topology and rooting, 58 and 56 of them remain clean—presence confined to one clade with no recovered absence inside it (Fig. 4c; Supplementary Fig. S8a). For this test, the 21 panel genomes absent from the GTDB trees were placed beside their closest relative on the project tree, so it is not fully independent of the project tree. Forty-five gains do not depend on where the root of the deep tree is placed, because the intron is confined to one family that also contains a recovered intron-free relative. Even under a deliberately extreme grouping rule that merges every gain sharing a locus or a branch into a single unit, seven separate units remain; Supplementary Fig. S8b shows the depth of every gain.

To measure how annotation affects this kind of inference, we also reconstructed gains from tRNAscan-SE intron calls alone, using the same loci and maximum-likelihood tree and an equally conservative rule (Methods). This annotation-based reconstruction yields 127 high-confidence gain calls (Supplementary Table S9). Checking their absences against the census shows how false absences arise (Supplementary Fig. S7). Allowing boundaries to differ by up to three nucleotides (the precision of the census), 53 of the 127 calls rested on at least one relative scored as intron-free that carries an intervening sequence at the same position (26 when the exact position is required), and for 34 of them that relative was the nearest one. Such insertions are often only partly similar in sequence to the called intron, so some may be independent insertions at the same site; either way the relative is not intron-free there. Of the 83 annotation-based calls that still satisfy the full gain rule once these false absences are recoded, 79 are census gains (Fig. 4d). Conversely, 47 census gains are absent from the annotation-based set; 18 of these involve sequences that tRNAscan-SE missed in at least one genome, and 14 of the 129 census gains consist only of missed sequences and are therefore invisible to annotation.

### 2.4 Intervening sequences sit at splicing-motif boundaries, including those the annotator missed

If introns persist only where they can be spliced out, they should sit at boundaries that can form a BHB motif far more often than chance. Two features of this test need care. First, the scoring rule matters. The relaxed BHB class, in which three of the four central pairs form and no flanking helix is required, is so permissive that some BHB-like class (canonical or relaxed) arises by chance at about two-thirds of arbitrary boundaries once the placement uncertainty of the census is allowed for (Fig. 5a). We therefore used the canonical classes—all four central pairs plus at least one flanking helix—as the test metric. Second, the test must use sequences found without the motif. Missed sequences are ideal for this, because the annotator, which searches for BHB-like structure, did not find them.

**Figure 5.**
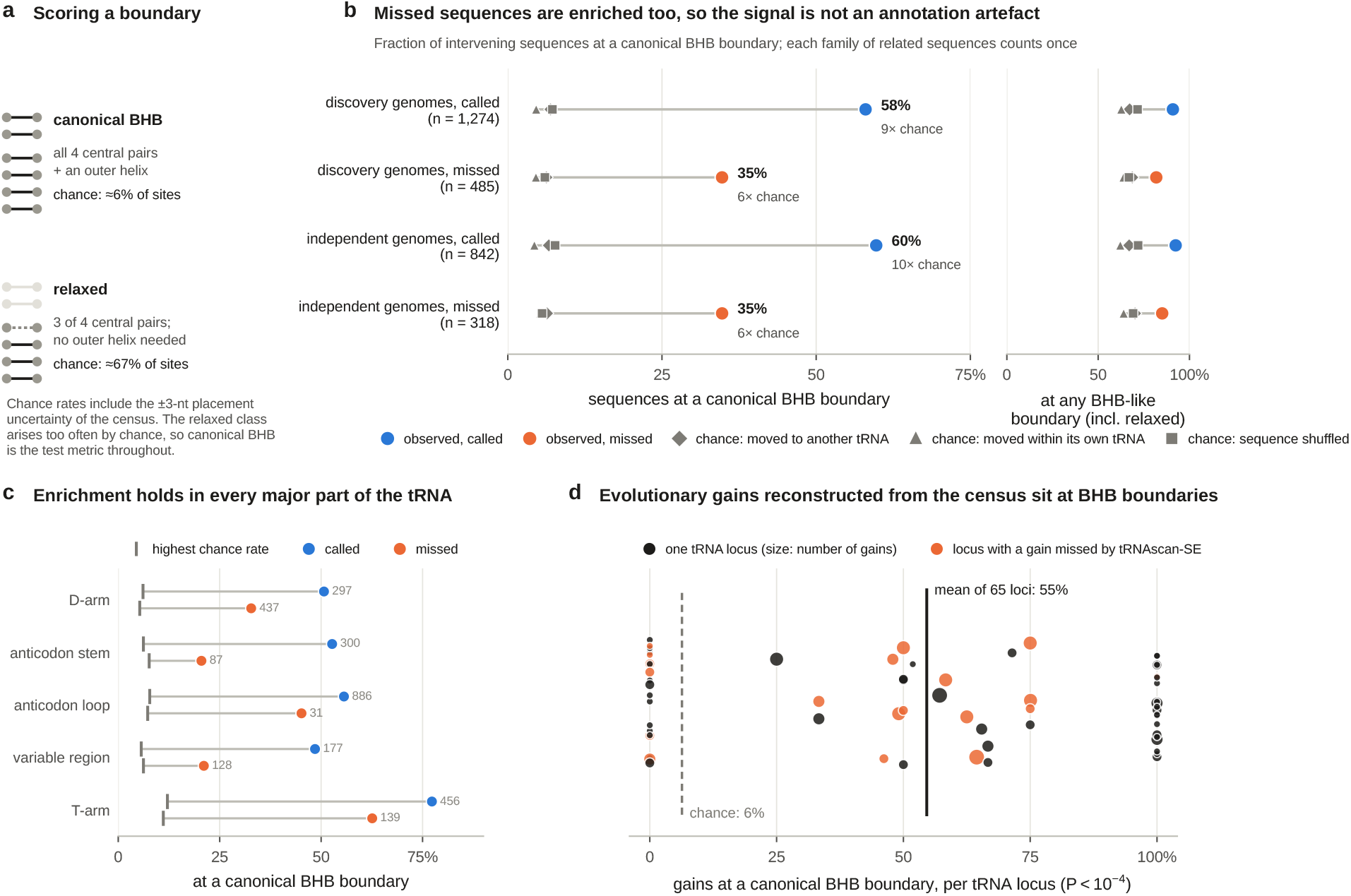
Intervening sequences sit at splicing-motif boundaries far more often than chance, including those the annotator missed. **a**, Scoring rules. A boundary is canonical BHB when all four central base pairs and at least one outer helix can form; the relaxed class requires only three central pairs and no outer helix. Chance rates are the average of the three null models for called introns in the discovery panel, with the *±*3-nucleotide placement window. **b**, Fraction of intervening sequences at canonical BHB boundaries (left) and at any BHB-like boundary including the relaxed class (right), for sequences called or missed by tRNAscan-SE in the discovery and independent panels. Coloured circles are observed values; grey symbols are the three null models. Each family of closely related sequences counts once. **c**, The same comparison by region of the tRNA (discovery and independent panels combined); grey ticks mark the highest of the three null rates, and numbers give sequence families. The few sequences in the acceptor stem and junctions are summarised as “other” in Supplementary Note 3; families containing both called and missed sequences contribute to both categories. **d**, Canonical BHB fraction of the 129 census gains, one circle per tRNA locus (area proportional to the number of gains; orange, loci with a gain missed by tRNAscan-SE). The solid line is the mean over loci and the dashed line the host-reassignment expectation.

We compared each intervening sequence with three null models that preserve different features of the data: the same sequence inserted at another tRNA’s insertion site, the same sequence moved at least eight nucleotides within its own tRNA, and a shuffled version of the sequence that keeps its dinucleotide composition. Every comparison used the same *±*3-nucleotide placement window as the observed data, and closely related sequences were collapsed into families so that near-identical copies counted once. In the discovery panel, 58% of called introns sat at a canonical BHB boundary, compared with 5–7% under the three nulls (Fig. 5b). Crucially, 35% of missed sequences did so as well, compared with 5–6% under the nulls (*P <* 10^*−*4^ for each comparison). The independent panel gave the same values (60% and 35%), and requiring the exact census boundary instead of a window lowered every fraction but preserved the contrast (called 37%, missed 19%, nulls 1–3%; Supplementary Fig. S3).

These results distinguish the biological signal from the annotation effect. Because missed sequences were found without any reference to the motif, their roughly six-fold enrichment (5.5-to 7.6-fold depending on the null model) cannot be produced by the annotator’s search. At the same time, the gap between called and missed sequences shows that the annotator preferentially finds the clearest motifs, so BHB fractions computed only from annotated introns overstate how strictly the motif is realised; with the relaxed class included, for example, 95% of the 127 annotation-based gain calls (98% when dependency units are weighted equally) appear BHB-compatible (Supplementary Note 3; Supplementary Table S10). The enrichment held in every major part of the tRNA (Fig. 5c; Supplementary Note 3). It was strongest for T-arm insertions, where even missed sequences formed canonical motifs 63% of the time, and weaker but still well above chance for D-arm (33%), anticodon-stem (20%) and variable-region (21%) insertions, where some missed sequences may use relaxed motifs or have imprecise boundaries.

The same pattern holds for the evolutionary gains reconstructed from the census. Weighting each of the 65 loci equally, 55% of gains sat at a canonical BHB boundary, compared with 6% after reassignment to another tRNA’s insertion site, 4% after moving within the same tRNA and 8% after shuffling (*P <* 10^*−*4^; Fig. 5d); 46 of the 65 loci lie above the host-reassignment expectation. Gains that involve missed sequences show the same direction (30% versus 6%; 19 loci, *P* = 0.007), as do the 14 gains made up only of missed sequences—the subset fully independent of annotation—although they are too few for a firm conclusion (26% versus 4%; 11 loci, *P* = 0.06). The annotation-based gain calls give a similar canonical result (72% of all 127 calls; 59% versus 8% when each of eight dependency units is weighted equally; Supplementary Note 3; Supplementary Table S9).

### 2.5 No single route of intron arrival explains the pattern

A splicing-motif requirement explains where introns can stay, not how they arrive. We tested the most direct general explanations, each by asking what pattern it predicts (Fig. 6).

**Figure 6.**
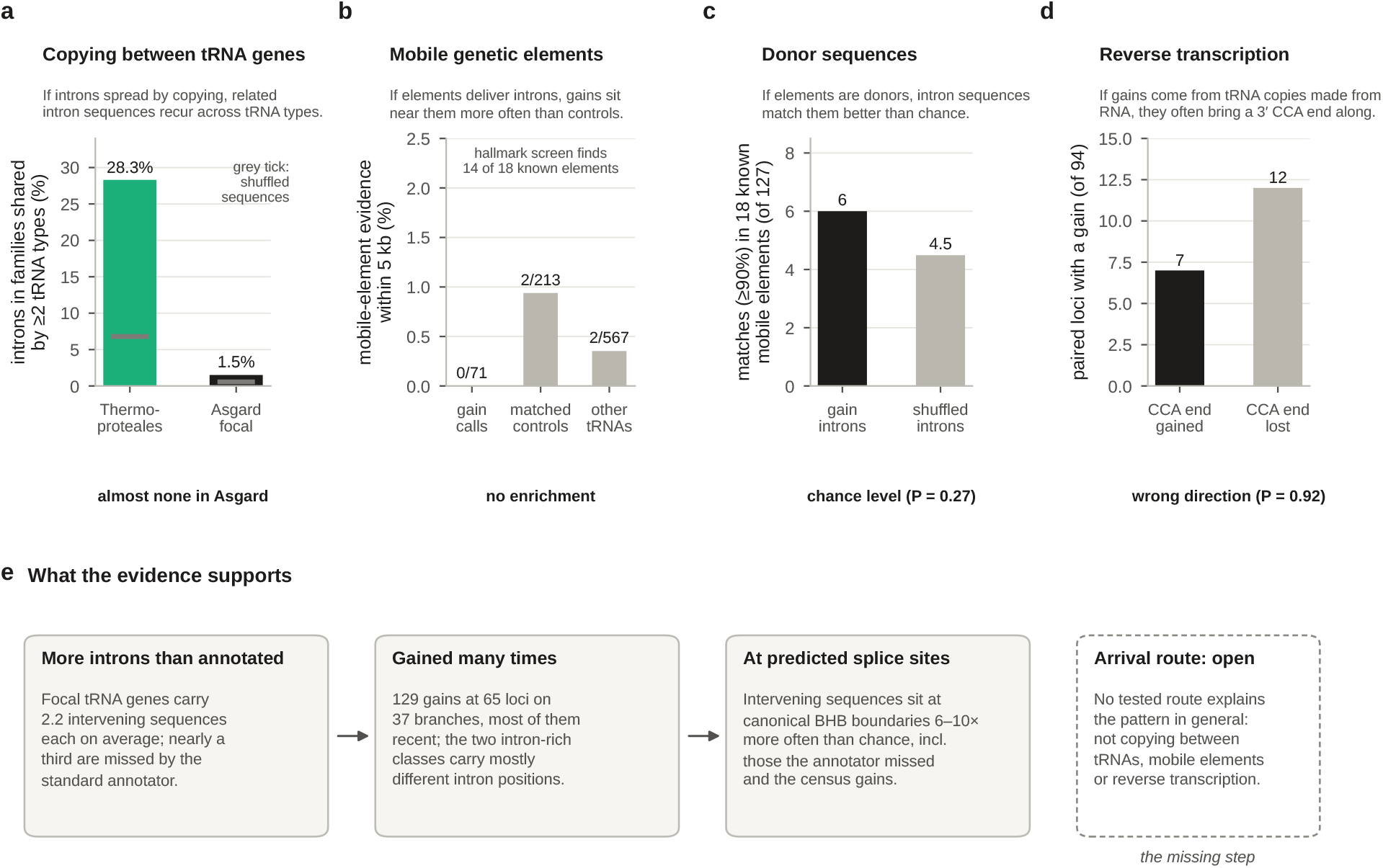
No tested route of intron arrival explains the pattern in general. Each panel states what the route predicts and shows the test. **a**, Copying between tRNA genes: percentage of introns, within each genome, that belong to families of *≥*80% identity spanning two or more tRNA types, computed identically for Thermoproteales (a positive control for intron transposition) and focal Asgard lineages from the full census; grey ticks, the same analysis on dinucleotide-shuffled sequences. **b**, Mobile genetic elements: percentage of annotation-based gain calls, matched control tRNA genes and other tRNA genes with strong mobile-element evidence within 5 kb. **c**, Donor sequences: gain introns matching a sequence in 18 published Asgard mobile elements at *≥*90% similarity, observed versus the mean for shuffled introns. **d**, Reverse transcription: gains accompanied by a newly encoded 3^*′*^ CCA end versus changes in the opposite direction. Panels **b**–**d** use the 127 annotation-based gain calls and do not cover the gains found only by the census. **e**, Summary of what the evidence supports and what remains open.

#### Copying between tRNA genes

If introns spread by copying themselves from one tRNA gene to another, as proposed for Thermoproteales (Sugahara et al., 2008; Fujishima et al., 2010), closely related intron sequences should occur in different kinds of tRNA within the same genome. Using the full census and the same clustering procedure for both groups (Supplementary Table S11), 28.3% of the 272 intervening sequences of eight Thermoproteales genomes belonged to families of at least 80% identity that span two or more tRNA types, against 6.8% for shuffled sequences. In 12 focal Asgard lineages the figure was 1.5% of 1,062 sequences, against 0.6% for shuffled sequences (Fig. 6a). At 90% identity the figures were 12.5% for Thermoproteales and 0.2% for Asgard (shuffled sequences 0% in both), whereas at 70% chance matches are common in both groups (Supplementary Fig. S4). A complementary analysis of published introns gave the same contrast: complete-linkage clustering at 70% identity found 26 recurrent families among 282 Thermoproteales introns but only one weak cross-isotype family among the 181 annotated introns of the three complete Asgard genomes (Supplementary Note 4). The host genes of that family were only 58–96% identical, so copying of an entire intron-containing tRNA gene does not explain it, and whole-gene duplication is not a general explanation either.

#### Mobile genetic elements

These tests were applied to the annotation-based gain calls (Supplementary Table S12). Mobile-element context was scored from hallmark-protein homology, exact mapping of published elements and local sequence features; hallmark-protein homology alone recovered 14 of 18 mobile elements published for these genomes (15 when the published geNomad calls are added). Strong mobile-element evidence lay within 5 kb of none of the 71 calls at well-assembled loci (57 host tRNA genes), compared with 2 of 213 matched control tRNA genes (three per call; 196 of them intron-free) and 2 of 567 other tRNA genes, and no window from 2 to 20 kb showed enrichment (Fig. 6b). Because mobile-element evidence is rare near any tRNA gene, this comparison could detect only a strong association. Six of 127 gain introns matched a sequence in the 18 published mobile elements at *≥*90% similarity, compared with 4.5 expected for shuffled introns (*P* = 0.27; Fig. 6c).

#### Reverse-transcribed RNA

A tRNA copy reverse-transcribed from RNA and inserted into the genome might bring a 3^*′*^ CCA end with it. Among 94 paired loci, a newly encoded CCA appeared together with a gain in 7 cases and changed in the opposite direction in 12 (*P* = 0.92 for the predicted direction; Fig. 6d). Only three credible reverse-transcriptase genes were present in the panel, none within 5 kb of a gain, and no gain combined the signatures expected of this route.

Negative results of this kind do not prove that a mechanism never operated: footprints decay, donors may be unsampled, and different gains may have had different origins. The mobile-element, donor and reverse-transcription tests used the annotation-based gain calls and therefore do not cover the 47 gains found only by the census. They do show that none of these routes provides a general explanation of the Asgard pattern (Fig. 6e).

### 2.6 The splicing enzyme is widespread but does not explain where introns accumulate

Processing so many introns requires a capable splicing endonuclease. An EndA catalytic-subunit homologue was found in 58 of the 60 discovery genomes; 44 contained two distinct full-length paralogues consistent with the two-subunit (*αβ*)_2_ architecture, and the remaining 16, including the two without a detectable homologue, could not be resolved, mostly because of assembly fragmentation. Known catalytic and substrate-recognition residues were broadly conserved, and burden was high in genomes of both architecture categories (Supplementary Fig. S6). Across the panel, EndA sequence divergence was strongly correlated with burden (*ρ* = *−*0.75), but this mostly reflects shared ancestry: within families, neither the length of a substrate-recognition loop nor sequence divergence was associated with annotated intron burden (minimum false-discovery-adjusted *P* = 0.112; Supplementary Table S13). Repeating the test with census-corrected burden gave an apparent association with divergence, but it was driven by 13 near-identical Freyrarchaeaceae assemblies; with near-identical genomes merged, neither feature was significant (loop length *ρ* = *−*0.52, *P* = 0.06; divergence *ρ* = *−*0.44, *P* = 0.11; unadjusted; 14 lineages in four families). With so few independent lineages a modest effect cannot be excluded, but EndA appears to be a broadly distributed, permissive background rather than the reason why particular lineages accumulated so many introns; its substrate range would have to be measured biochemically.

## 3 Discussion

Asgard tRNA genes are even more heavily interrupted than the first complete genomes suggested (Valentin-Alvarado et al., 2024). In the three focal families nearly every tRNA gene carries an intervening sequence, about three-quarters carry at least two, and the average gene carries more than two—well above *Pyrobaculum*, the most intron-rich reference archaeon in our comparison. The extra sequences had been hidden because the standard annotator systematically misses insertions in the D-arm. With the corrected catalogue, the evolutionary picture is consistent: introns were gained many times, on many branches and apparently independently in two classes, and they sit at boundaries that can form the canonical splicing motif far more often than chance. What remains unknown is how intron DNA enters a tRNA gene.

The annotation gap has consequences beyond this study. tRNAscan-SE missed almost nothing in our reference archaea, including intron-rich Thermoproteales, but Asgard introns occupy positions and combinations that are rare elsewhere. Curators of the first complete Asgard genomes had already noticed introns that the software did not report (Valentin-Alvarado et al., 2024); our results show that the gap is systematic, concentrated in the D-arm and largest in the most intron-rich lineages. Its omissions had effects that pointed in opposite directions. They removed introns from the census, making the phenotype look less extreme, and they inserted false absences into the evolutionary data, making some gains appear that may not be real. A third effect is subtler: because the annotator finds the clearest motifs most easily, structural statistics computed from its calls overstate how strictly the motif is realised. Each of these biases is invisible from inside an annotation-based analysis and becomes measurable only with an independent census. Simple homology-based checks like the one used here could be applied wherever annotation software is used to study the evolution of the very feature it was designed to detect. Interestingly, the missed set also includes many T-arm insertions with strong canonical motifs, so missing an intron is not simply a matter of motif weakness; it may also reflect how the software aligns long, multiply interrupted precursors to its models.

The historical conclusion rests on a reconstruction whose intron states do not depend on annotation, and on several independent lines within it. Gains fall on three dozen branches, most of them terminal; the internal gains remain consistent with the GTDB reference phylogenies; many do not depend on the root; and the two intron-rich classes, which are not sister lineages, carry mostly different intron positions. The exact number of gain events remains uncertain—draft genomes leave gaps, and conservative grouping trades resolution for safety—but even the most conservative grouping leaves seven separate units, and most analyses imply many more. Losses probably occurred as well, although parsimony cannot fully separate them from repeated gains, and the frequency of recent gains and of positions that changed state repeatedly suggests continual turnover rather than a fixed set of ancient introns. Repeated use of the same positions in different lineages is what one would expect if only a limited set of positions in each tRNA can host a removable intron, linking the historical result to the structural one.

Tests of the BHB association that use annotated introns are partly guaranteed to find it, because the annotator itself searches for BHB-like structure. The missed sequences remove that guarantee, and they still show a roughly six-fold enrichment for canonical motifs; the census gains show a roughly nine-fold enrichment. This supports the view that the predicted ability to be spliced constrains where introns persist. Our data cannot tell whether the constraint acts at insertion, through a bias in where new sequence lands, or afterwards, through the loss of insertions that disrupt splicing; the two are not mutually exclusive. Nor is a canonical motif strictly required: a substantial minority of sequences, particularly in the D-arm, lack one. Some of these may be removed through relaxed motifs, which archaeal EndA enzymes with broad substrate specificity can cleave (Randau et al., 2005; Hirata et al., 2012); others may reflect imprecise boundaries or insertions that are not introns at all.

The comparison with Thermoproteales is instructive. Both groups carry many noncanonical introns and both depend on BHB-mediated splicing, yet in Thermoproteales a large fraction of introns belong to families shared across tRNA types, consistent with transposition (Fujishima et al., 2010), whereas in Asgard almost none do, even in the complete census. The same constraint on the RNA product can therefore accompany very different histories of DNA arrival. The Asgard pattern is not simply a scaled-up version of the Thermoproteales one.

Our tests of arrival routes were designed to detect a general explanation and found none. Copying between tRNA genes, mobile elements and reverse transcription each predict signatures that were tested directly and found absent or not enriched, although the mobile-element test had limited power and none of these tests yet covers the gains found only by the census. These results leave room for routes that leave few genomic traces, for routes that operated too long ago for traces to survive, and for a mixture of routes. The abundance of recent, lineage-specific gains, and the availability of closely related genomes that differ in individual introns, make Asgard tRNA genes a promising system in which to catch insertion in progress, for example through population-level sequencing of natural communities.

Several limitations apply. Most genomes are metagenome-assembled drafts, which can hide or fragment loci; we therefore scored absences only where a locus was recovered and used draft genomes mainly for presence. The census inspects only tRNA genes that tRNAscan-SE detected, so a gene disrupted beyond recognition would be invisible to both methods, and insertions within a few nucleotides of either end of a gene are detected less reliably. Census boundaries are precise to a few nucleotides, which is why every structural test applied the same placement window to observed data and nulls. The BHB classifier encodes published sequence rules and predicts the ability to form a motif, not cleavage efficiency. Finally, all analyses are comparative. None of the newly identified intervening sequences has yet been shown to be spliced in an Asgard cell; sequencing of precursor and mature tRNAs from the same environments (Valentin-Alvarado et al., 2024), or cleavage assays with Asgard EndA, would test this directly.

In summary, the heavily interrupted tRNA genes of these Asgard archaea carry more introns than standard annotation reports, were built by many separate gains in at least two lineages, and retain their introns where the splicing machinery can act. Finding out how the intron sequences arrive is now the central open problem.

## 4 Materials and Methods

### 4.1 Genome panels

The discovery panel comprised 60 Asgard genomes: the three complete wetland-soil genomes (Valentin-Alvarado et al., 2024), dense sampling of the focal families, three near-basal comparator genomes, and one or more representatives of other families for context. Taxonomy followed GTDB releases 226 and 232 (Parks et al., 2020, 2022). The independent panel comprised 30 public assemblies not used for discovery, selected before any tRNA analysis from assembly level, completeness, contamination, contiguity, GTDB taxonomy and representative status, with four non-focal genomes chosen for proximity to the focal families on the species tree; intron burden and motif content were not selection variables. Eighteen non-Asgard archaeal genomes, including eight Thermoproteales, served as references. For descriptive summaries, discovery genomes separated by less than 0.005 substitutions per site on the species tree were merged by single linkage into 41 lineages; the largest merged group contained 13 near-identical Freyrarchaeaceae assemblies. Accessions and quality metrics are listed in Supplementary Table S1.

### 4.2 tRNA annotation

tRNAs were predicted with tRNAscan-SE 2.0.12 in archaeal mode (Chan et al., 2021); predicted pseudogenes were excluded from burden estimates. Coordinates, strand, intron boundaries and mature sequences were validated against the genome sequence, and ARAGORN 1.2.41 provided an independent check of gene calls and anticodon-loop introns (Laslett and Canback, 2004). The “mature” sequence of a gene was its sequence after removal of every tRNAscan-SE intron (Supplementary Table S2).

### 4.3 Caller-independent census of intervening sequences

Compact reference tRNAs were mature sequences, with any genomically encoded 3^*′*^ CCA removed, of 70–78 nucleotides for class I isotypes or 80–89 nucleotides for leucine and serine tRNAs, pooled from all 108 genomes (Supplementary Table S1), excluding predicted pseudogenes, and de-duplicated. For each tRNA gene, the full precursor sequence (3^*′*^ CCA removed) was compared with the 12 compact references of the same isotype and anticodon that shared the most 5-mers with it (references of the same isotype were added when fewer than three matched the anticodon; genes of undetermined isotype and selenocysteine tRNAs were compared with all references), excluding the gene itself. Each pair was aligned with an affine-gap semi-global algorithm (match +3; mismatch *−*2; gap in the reference, that is an insertion in the gene, opened at *−*8 and extended at *−*0.25 per nucleotide; gap in the gene opened at *−*8 and extended at *−*2; end gaps free), and the highest-scoring alignment was retained. Every internal run of *≥*10 gene nucleotides aligned to reference gaps was recorded as an intervening sequence; runs at the alignment ends were ignored. An intervening sequence whose overlap with a tRNAscan-SE intron covered at least half of the shorter of the two was labelled *called*; all others were labelled *missed*. The fully spliced sequence of each gene was obtained by removing every intervening sequence. A sensitivity analysis used a 13-nucleotide threshold.

To locate intervening sequences on the cloverleaf, each boundary was projected through the alignment onto the reference tRNA and assigned to a structural element and an approximate Sprinzl position (Sprinzl et al., 1998) using the secondary structure reported by tRNAscan-SE for that reference.

### 4.4 Independent checks of the census

Six checks were made. (i) Recovery of tRNAscan-SE introns and agreement of boundaries. (ii) The rate of extra insertions in the 18 reference genomes. (iii) Per-gene agreement with the manually curated intron counts of the three complete genomes, obtained by comparing the published R2DT-curated and tRNAscan-SE counts (Valentin-Alvarado et al., 2024). (iv) Agreement with a covariance-model alignment: the “mature” sequences of the discovery and independent panels were aligned to the archaeal tRNA covariance model with Infernal 1.1.5 cmalign (Nawrocki and Eddy, 2013), and we counted, for each gene, runs of *≥*10 residues in insert or local-end states between consecutive consensus columns. (v) Simulation: real intervening sequences of *≤*120 nucleotides drawn at random from the census were inserted at random internal positions of 1,212 distinct compact tRNA genes with no detected insertion (one insertion per gene, and separately two), and the census was rerun with the host’s own sequence and identical copies removed from the reference pool; an insertion was recovered when a detected run overlapped at least half of it. (vi) A model-free length check of “mature” class I tRNAs. Results are in Supplementary Table S5 and Supplementary Note 1.

### 4.5 Burden estimates and the independent-panel test

Burden was the number of intervening sequences per non-pseudogene tRNA gene. Discovery-panel summaries average genomes within merged lineages and then lineages within groups. For the independent panel, burden was averaged over the genomes of each family. The test statistic was the difference between the median of the two focal families and the median of the 19 others. Because only 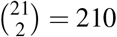 assignments of the focal label are possible, the null distribution was enumerated exactly. Associations of the missed fraction with annotated burden and assembly metrics used Spearman correlation across genomes; because related genomes are not independent, these are descriptive.

### 4.6 Census-based intron positions and gain reconstruction

Corresponding tRNA loci across genomes were defined by combining mature-sequence similarity, covariance-model coordinates and conserved neighbouring protein orthogroups (MMseqs2 18.8cc5c; Steinegger and Söding, 2017); loci supported by synteny (Tier 1) or by sequence and structure (Tier 2) were used. Within each locus, one reference member was chosen (a genome of high or moderate tRNA-repertoire quality whose spliced length was closest to the locus median), and every member’s intervening sequences were mapped into its coordinates by pairwise alignment of fully spliced sequences. Mapped positions were sorted and grouped into intron positions (characters), a new character starting whenever a position lay more than three nucleotides from the first position of the current one. Each genome was scored 1 if it had an insertion at the position; 0 if its copy of the locus was recovered with no insertion within six nucleotides; and unknown if the locus was not recovered or an insertion belonging to another character lay within six nucleotides of the character’s median position. No tRNAscan-SE intron call was used to assign states (Supplementary Table S7).

The species tree was inferred from the 53 GTDB archaeal marker proteins, taken from the GTDB release 226 per-marker alignments, subset to the 60 discovery genomes, stripped of columns with more than 50% gaps and concatenated (12,891 columns), with IQ-TREE 2.4.0 (model selected by ModelFinder under BIC from LG, WAG and JTT with model-defined or empirical frequencies and E, I, G4 or R4 rate heterogeneity, giving LG+F+R4; 1,000 ultrafast bootstrap replicates; Minh et al., 2020; Kalyaanamoorthy et al., 2017; Hoang et al., 2018), and midpoint-rooted, with bootstrap support transferred to the rooted tree by bipartition matching. A high-confidence gain required that presence be confined to one clade with no recovered absence inside it (unknowns allowed), support *≥*80 for that clade (or for the clade containing a single genome and its sister), at least one recovered absence in a genome of high or moderate tRNA-repertoire quality, at least one present copy classed as a high-confidence tRNA, and at least one present copy at a Tier 1 locus; adding Tier 2 loci gave 173 gains at 88 loci on 43 branches. Intron positions shared across the focal families were not excluded, so the location of deep gains could be tested rather than assumed. A gain was root-independent when presence was confined to one family that also contained a recovered absence. Topological robustness was assessed on the project tree with branches of support *<*95 collapsed and on the GTDB release 226 and 232 reference trees, pruned to the panel, with the 21 panel genomes absent from these trees (non-representative genomes of GTDB species) grafted beside their closest relative on the project tree. Every position was also scored by Fitch parsimony on the rooted tree, treating unknown states as wildcards (Fitch, 1971). Positional sharing between the two intron-rich classes was summarised at loci recovered in at least one genome of each class.

### 4.7 Annotation-based gain calls and the census check of absences

For comparison, gains were also reconstructed from tRNAscan-SE intron calls alone. Each called intron was assigned to its locus and to a column of the Infernal structural alignment of mature sequences; a locus–column pair defined a character, scored present where a genome’s copy carried a called intron at that column, absent where the copy was recovered without one, and unknown otherwise. A high-confidence call required a Tier 1 locus, a high-confidence host tRNA, presence confined to one clade with no recovered absence inside it, at least one present and one absent genome of high or moderate tRNA-repertoire quality, and support *≥*80 for the presence clade (or for the parent node of a single genome). Clades for this rule were defined on the same maximum-likelihood tree as written by IQ-TREE, which places the root arbitrarily at the first taxon (here the Asgardarchaeaceae genome GCA_040225955.1), so clade membership for this rule depends on that arbitrary root; census gains used the midpoint-rooted tree and were additionally tested on three alternative trees. Characters present in both Atabeyarchaeaceae and Jordiarchaeaceae with presence inferred at their common ancestor, or whose distribution across the focal families could not be classified, were excluded, so this rule detects recent gains only. Calls were grouped into dependency units by joining calls that share a locus, a locus and alignment column, or the same branch. Topological stability was assessed on the project tree, the GTDB release 232 tree, the family-level backbone of Zhang et al. (2025) and the GTDB release 226-based alternative of Manzano-Morales and Gabaldón (2026). A call was root-independent when presence was confined to one family and an intron-free relative from that family, supported at *≥*80, sat beside the presence clade.

To check absences, the intron position of each call was mapped into the coordinates of a reference member of the locus, and every genome scored as absent was searched for a census-detected intervening sequence within 0, *±*1 or *±*3 nucleotides of that position. Such genomes were recoded as present and the full gain rule, including the support of the enlarged presence clade, was re-applied. Calls were matched to census positions through the insertion that overlapped each call’s representative intron. Further details are given in Supplementary Note 2.

### 4.8 BHB classification and null models

The classifier encodes published motif definitions (Marck and Grosjean, 2003): a four-base-pair central helix (Watson–Crick or G–U pairs) formed between the four exon nucleotides upstream of the 5^*′*^ bulge and the corresponding intron nucleotides upstream of the 3^*′*^ bulge, three-nucleotide bulges containing the cut sites, and flanking helices of at least two contiguous pairs formed by exon (h) or intron (h^*′*^) sequence. Classes hBHBh^*′*^, hBH and HBh^*′*^ were scored as canonical; a central helix with three of four pairs and no outer-helix requirement was scored as relaxed. Because the outer-helix requirement applies only to boundaries with all four central pairs, a boundary with four central pairs but no outer helix is scored as not BHB-like, whereas one with three central pairs is scored as relaxed; canonical fractions, the test metric, are unaffected, but the any-class fractions should be read with this rule in mind. On 24 published junctions and 12 composition-preserving designed negatives, every positive was classed as compatible (20 with exactly the published class and 4 as relaxed) and every negative as incompatible (Supplementary Fig. S5). The classifier predicts the potential to form a motif, not cleavage.

Each intervening sequence was evaluated in the fully spliced context of its gene. Census boundaries were allowed to shift by up to *±*3 nucleotides (excised length fixed), and the best class within this window was recorded; the identical window was applied to every null. Nulls were (i) the sequence inserted at the insertion site of a randomly chosen other intervening sequence (host reassignment), (ii) the sequence moved to a random boundary at least eight nucleotides from its own within the same gene, and (iii) a dinucleotide-preserving shuffle, with 10 replicates per sequence and null. Identical sequence–host pairs were counted once, and sequences of the same isotype were collapsed into families by greedy clustering against family representatives (identity *≥*75%, lengths within 30%); family values were averaged. Families containing both called and missed sequences were excluded from the called-versus-missed comparison and, in the regional breakdown, contribute to both categories. One-sided *P* values compared the observed canonical fraction with 20,000 simulated draws from the per-family probabilities of the most permissive of the three nulls. For census gains, the values of each gain’s member insertions were averaged, gains were averaged within loci, and loci were weighted equally, with *P* values computed in the same way. For the annotation-based calls, the tRNAscan-SE boundary was evaluated exactly, introns were reassigned among gain hosts within strata of intron position (canonical 37/38 or noncanonical) and intron length (2,000 permutations), the position null moved each intron at least three nucleotides, and statistics were averaged within dependency units and then across units, with *P* = (*b* + 1)*/*(*B* + 1) (Phipson and Smyth, 2010).

### 4.9 Tests of arrival routes and EndA

Sequence reuse was assessed within each genome by single-linkage clustering of intervening sequences with pairwise identity at or above 70%, 80% or 90% (lengths within 20%), counting sequences in families that span at least two tRNA isotypes; dinucleotide-shuffled sequences provided the null, and one genome per merged lineage was used for the focal families. Mobile-element analyses, applied to the annotation-based gain calls, combined hallmark-protein homology (proteins predicted with Prodigal 2.6.3), exact mapping of 18 published Atabey mobile elements (whose boundaries came from the published geNomad 1.5.0 output; Camargo et al., 2024), local composition and repeat features, three same-genome control tRNA genes per call matched for isotype, anticodon, gene density and distance to the contig end (preferentially intron-free), and a donor search of gain introns against the published elements with a mononucleotide-shuffle null. The RNA/cDNA analysis combined the direction of encoded-CCA change between paired loci, local partial-tRNA fragments and the location of credible reverse-transcriptase genes. EndA homologues were identified against structural *α* and *β* references (query coverage *≥*0.60, *E ≤* 10^*−*8^), and associations with burden were tested by Spearman correlation after centring within GTDB family (families with *≥*4 genomes), with Benjamini–Hochberg correction (Benjamini and Hochberg, 1995); the census-based repeat used intervening sequences per gene, both for genomes and for merged lineages (families with *≥*2 lineages). Details are given in the Supplementary Information.

### 4.10 Software and reproducibility

tRNA prediction, locus matching, tree inference, annotation-based gain calls, mobile-element and reverse-transcription screens and the EndA inventory used Python 3.12, tRNAscan-SE 2.0.12, ARAGORN 1.2.41, Infernal 1.1.5, IQ-TREE 2.4.0, MMseqs2 18.8cc5c, Prodigal 2.6.3 (Hyatt et al., 2010), Biopython 1.85 and RapidFuzz 3.14; all package versions are recorded in the environment lock files of the archive. The census and all downstream analyses used Python 3.13 with NumPy 2.5, pandas 3.0, SciPy 1.18, NetworkX 3.6 and Matplotlib 3.11; the census alignment is a short C routine included in the archive. Starting from the outputs of the upstream steps, one command regenerates every number in the paper, every figure and Supplementary Tables S1–S14 in about five minutes on a laptop; all random procedures use recorded seeds, and Supplementary Table S14 lists checksums of every input, code and result file (Supplementary Note 5).

## Data availability

All genome assemblies are public NCBI accessions listed in Supplementary Table S1. Genome sequences used, all intermediate and result files, source code, environment specifications, checksums and step-by-step reproduction instructions are deposited at Zenodo (https://doi.org/10.5281/zenodo.23013520; version accompanying this article). Third-party sequences remain credited to their original repositories.

## Code availability

The analysis code, including the census, its independent checks, the census-based gain reconstruction, the structural tests and the figure scripts, is available at the Zenodo record above.

## Acknowledgements

We acknowledge the open-source data resources, public databases, software and data-sharing practices that made this unfunded study possible. S.G. and A.K.G. acknowledge the Department of Zoology, Siksha Bhavana, Visva-Bharati, where they are M.Sc. students.

## Author contributions

K.S. contributed to all aspects of the study, including conceptualisation, methodology, software, validation, analysis, investigation, data curation, visualisation, supervision, project administration, and writing (original draft, review and editing). S.G. and A.K.G. performed the study work and analyses, contributed to writing (original draft), and reviewed and edited the manuscript.

## AI disclosure

Artificial intelligence (AI) tools assisted with workflow design, code generation, data analysis, figure preparation and language editing. No AI tool is listed as an author. The authors reviewed and verified the code, analyses and outputs and take full responsibility for the manuscript and its conclusions.

