## Supplementary information for "Hidden introns and repeated gains shape the heavily interrupted tRNA genes of Asgard archaea"

#### Contents

**Supplementary Note 1. Caller-independent census** **1**

**Supplementary Note 2. Gain reconstruction** **3**

**Supplementary Note 3. Structural tests** **3**

**Supplementary Note 4. Independent panel, sequence reuse, mobile elements and EndA** **6**

**Supplementary Note 5. Reproducibility** **7**

**Supplementary Figures** **8**

**Supplementary Tables** **12**

#### Supplementary Note 1. Caller-independent census

**Procedure.** Compact references were mature tRNA sequences (genomic 3′ CCA removed) of 70–78 nt (class I) or 80–89 nt (Leu, Ser), excluding predicted pseudogenes, pooled from the 60 discovery, 30 independent and 18 reference genomes (108 genomes) and de-duplicated. Each tRNA gene's full precursor sequence was compared with up to 12 compact references of the same isotype and anticodon (ranked by shared 5-mers), and the best-scoring affine-gap semi-global alignment was retained (match $+3$, mismatch $-2$; insertion in the gene: open $-8$, extend $-0.25$; deletion in the gene: open $-8$, extend $-2$; free end gaps). Internal gene-only runs of $\geq$10 nt were recorded as intervening sequences. Overlap of at least half of a tRNAscan-SE intron defined *called*; all others were *missed*.

**Six independent checks.** (i) *Recovery of annotated introns:* 2,781 of 2,879 discovery-panel introns (96.6%) and 1,085 of 1,109 independent-panel introns (97.8%) were re-found; boundaries agreed exactly for 58% and within $\pm$3 nt for 88% (Figure S1d). (ii) *Reference archaea:* the census found 8 intervening sequences in 826 tRNA genes of 18 reference archaea, including 4 beside the 277 annotated introns of eight Thermoproteales genomes. (iii) *Expert curation:* of 118 tRNA genes of the three complete genomes matched to the published annotation, curators had added introns to 35 (adding 43 introns in total); the census found extra sequence in 33 of these, and found none in 74 of the 83 genes to which curators added nothing. (iv) *Covariance-model alignment:* for 3,698 genes of the discovery and independent panels, the Infernal alignment of "mature" sequences and the census agreed on the presence of a missed insertion for 97.4% of genes (both 1,259, neither

2,343, census only 29, Infernal only 67), agreed on the exact number for 94.4%, and the inserted lengths correlated with Pearson $r = 0.95$ (Figure S1b). (v) *Simulation:* with one real intervening sequence inserted at a random internal position of each of 1,212 compact genes, the census recovered 91.8%, placed 59% at the exact boundary and 92% within $\pm 3$ nt, and reported 0.07 spurious extra runs per gene; with two insertions per gene, recovery was 90.8% (Figure S1c). Recovery was lowest for insertions in the acceptor stem, within a few nucleotides of the gene ends, where free end gaps absorb insertions; such introns are rare. (vi) *Model-free length check:* after removal of annotated introns, 48% of focal-family class I genes were $\geq 88$ nt, against 16% in other Asgard families and 0.5% in reference archaea; after removal of all census insertions, the median length was 75 nt in the focal families (Figure S1a).

**Location of missed sequences.** Table N1 gives the location of every discovery-panel intervening sequence. The canonical position is almost never missed; D-arm insertions dominate the missed set. The table counts census runs: in 57 genes an annotated intron was split by the census into two or three runs at internal mismatches (65 extra runs), so the number of called runs (2,846) slightly exceeds the number of annotated introns re-found (2,781).

**Table N1.** Location of discovery-panel intervening sequences on the tRNA, by annotation status (all sequences in non-pseudogene tRNA genes).

| Region | Called | Missed | Missed (%) |
|---|---:|---:|---:|
| acceptor stem | 25 | 2 | 7% |
| junctions (8–9, 26) | 37 | 4 | 10% |
| D-stem | 107 | 188 | 64% |
| D-loop | 350 | 484 | 58% |
| anticodon stem | 394 | 122 | 24% |
| anticodon loop (non-canonical) | 339 | 40 | 11% |
| canonical 37/38 | 521 | 14 | 3% |
| variable loop | 126 | 74 | 37% |
| variable arm (class II) | 122 | 64 | 34% |
| T-stem | 187 | 62 | 25% |
| T-loop | 616 | 154 | 20% |
| not assigned (no structure for reference) | 22 | 27 | 55% |
| Total | 2,846 | 1,235 | 30% |

**Robustness.** Raising the minimum length to 13 nt reduced the number of missed sequences by about 10% and left every qualitative result unchanged (Figure S2a). Missed and called sequences have the same length distribution (Figure S2b). Across 90 Asgard genomes the missed fraction rises with annotated burden (Spearman $\rho = 0.47$, $P = 3 \times 10^{-6}$) but not with contig number ($\rho = 0.018$), completeness ($\rho = 0.13$) or N50 ($\rho = 0.0030$) (Figure S2c). Because near-identical genomes are not independent, we repeated the correlations on 71 units (discovery genomes merged into lineages, independent-panel genomes kept separate): annotated burden $\rho = 0.37$, $P = 0.001$; contig number $\rho = 0.10$; completeness $\rho = -0.07$; N50 $\rho = -0.04$.

**Limitations.** The census inspects only tRNA genes that tRNAscan-SE detected; a gene so disrupted that it was not called at all is invisible. Alignment boundaries are imprecise by a few nucleotides at repeats and mismatches. An intervening sequence is a candidate intron; its splicing has not been shown experimentally.

#### Supplementary Note 2. Gain reconstruction

**Character matrix.** The 193 multi-genome Tier 1 (syntenic) and Tier 2 (sequence and structure) loci contain 541 census intron positions in 172 loci (Supplementary Table S7); 163 positions include at least one missed insertion and 110 consist only of missed insertions. Of the 953 missed insertions in these loci, 305 lie at a position where the insertion was called in another genome, showing that annotation is often inconsistent between orthologous copies.

**Gains.** The primary rule (main text, Methods) yielded 129 high-confidence gains at 65 loci on 37 branches; 71 were on terminal branches, 45 were root-independent and 23 involved at least one missed insertion (14 only missed insertions). Adding Tier 2 loci gave 173 gains at 88 loci on 43 branches. On four trees—the project tree, the project tree with branches of support <95 collapsed, and GTDB releases 226 and 232—127 gains remained clean (presence confined to one clade with no recovered absence inside it) on all four and the remaining 2 on three (Figure S8a). This test is informative only for internal gains: terminal gains are clean on any tree, and every gain clade has support ≥95, so collapsing branches of support <95 cannot affect them. Of the 58 internal gains, 58 are clean on the GTDB release 226 tree and 56 on the release 232 tree. The 21 panel genomes absent from the GTDB trees were grafted beside their closest relative on the project tree, so this test is not fully independent of the project tree. Merging every gain that shares a locus or a branch leaves 7 units; merging only gains that share a locus leaves 65.

**Ancestral burst.** The two intron-rich classes, Jordiarchaeia and Atabeyarchaeia, are not sister groups: their common ancestor is also the ancestor of 14 other families in the panel on the project tree and of 27 and 26 other families on the GTDB release 226 and 232 trees. At the 18 loci recovered in both classes, 45 intron positions occur only in Jordiarchaeia, 13 only in Atabeyarchaeia and 22 in both; every shared position requires at least two changes under Fitch parsimony. The stem branches of the two classes carry 8 and 6 gains, respectively. Of all 541 positions, 104 require at least two changes.

**Annotation-based gain calls.** Gains reconstructed from tRNAscan-SE intron calls alone (main text, Methods) comprise 127 high-confidence calls (Supplementary Table S9). Because that rule excludes intron positions present in both Atabeyarchaeaceae and Jordiarchaeaceae with presence inferred at their common ancestor, it can detect recent gains only and cannot be used to test for an ancestral burst; the census-based rule has no such exclusion. Of the 127 calls, 49 are gains on all four topologies tested and 75 on most, and 51 are root-independent. Grouping calls that share a locus, a locus and alignment column, or a branch gives 8 dependency units; one unit contains 112 calls, so the grouping is deliberately severe. For each call, the intron boundary was mapped into the coordinates of a reference member of its locus, and every genome scored as absent was checked for a census-detected intervening sequence at the same mapped position (tolerance 0, ±1 or ±3 nt). Such genomes were recoded as present and the full rule re-applied, including support ≥80 for the enlarged presence clade (Table N2; Figure S7). Clades for the annotation-based rule are defined on the maximum-likelihood tree as written by IQ-TREE, which places the root arbitrarily at the first taxon (GCA_040225955.1, Asgardarchaeaceae); clade membership for this rule therefore depends on an arbitrary root. Census gains use the midpoint-rooted tree. Of the 83 calls without false absences at ±3 nt, 79 are census gains; overall 82 of the 127 calls are census gains, and 47 census gains are not among the 127.

#### Supplementary Note 3. Structural tests

**Classifier.** The classifier enumerates the four-pair central helix formed between exon positions −6 to −3 upstream of the 5′ cut and the corresponding intron positions upstream of the 3′ cut, three-nucleotide bulges,

**Table N2.** Census check of the absences behind the 127 annotation-based gain calls. "Calls with false absence": calls with at least one genome scored absent that carries an intervening sequence at the same position. "Nearest relative affected": the nearest intron-free relative used to support the call is among them. The two cause columns count genome–call pairs: a missed insertion, or an annotated intron assigned to a neighbouring alignment column. Remaining calls still satisfy the gain rule after recoding; dependency units use the grouping rule above. Units increase as calls are removed because links between otherwise unrelated calls are broken.

| Tolerance | Calls with false absence | Nearest relative affected | Cause: missed | Cause: other column | Remaining calls | Root-independent | Dependency units |
|---|---|---|---|---|---|---|---|
| annotation only | – | – | – | – | 127 | 51 | 8 |
| exact position | 26 | 19 | 63 | 42 | 107 | 39 | 11 |
| ±1 nt | 38 | 25 | 106 | 60 | 98 | 37 | 14 |
| ±3 nt | 53 | 34 | 127 | 170 | 83 | 31 | 16 |

and outer helices of 2–7 contiguous Watson–Crick or G–U pairs formed by exon (h) or intron (h′) sequence. Canonical classes require four central pairs and at least one outer helix; the relaxed class requires three central pairs and no outer helix. A boundary with four central pairs but no outer helix is therefore scored as not BHB-like, whereas one with three central pairs is relaxed; this affects only the any-class (canonical or relaxed) fractions, not the canonical test metric. On 24 published junctions and 12 designed negatives, all positives were compatible (20 with exactly the published class, 4 assigned to the relaxed class) and all negatives incompatible (Figure S5).

**All intervening sequences.** Table N3 gives observed and null fractions for sequence families with the ±3-nt placement window. Families were built by greedy clustering: each sequence joined the first family representative of the same isotype with ≥75% identity and length within 30%, otherwise it founded a new family. Families with both called and missed members (20 in the discovery and 14 in the independent panel) are omitted from the called and missed rows; Table N4 repeats the analysis at the exact census boundary (Figure S3). All one-sided $P$ values for canonical BHB are below $10^{-4}$. Table N5 breaks the comparison down by region; there, values are averaged within status and family, so a family with both called and missed members contributes to both rows.

**Table N3.** Canonical and any-class BHB fractions for intervening-sequence families (±3-nt window). Nulls: host = placed at another sequence's insertion site; position = moved ≥8 nt within its own gene; dinuc. = dinucleotide-preserving shuffle.

| | | Canonical BHB | | | | Any BHB-like | |
|---|---|---|---|---|---|---|---|
| Set | Families | Observed | Host | Position | Dinuc. | Observed | Nulls |
| Discovery, called | 1,274 | 0.58 | 0.069 | 0.046 | 0.072 | 0.91 | 0.63–0.72 |
| Discovery, missed | 485 | 0.35 | 0.063 | 0.046 | 0.060 | 0.82 | 0.64–0.69 |
| Independent, called | 842 | 0.60 | 0.067 | 0.043 | 0.077 | 0.92 | 0.62–0.72 |
| Independent, missed | 318 | 0.35 | 0.064 | 0.053 | 0.056 | 0.85 | 0.64–0.70 |

**Table N4.** As Table N3 but at the exact census boundary (discovery panel; families as in Table N3).

| | Canonical BHB | | | | Any BHB-like | |
|---|---|---|---|---|---|---|
| Set | Observed | Host | Position | Dinuc. | Observed | Nulls |
| Discovery, called | 0.37 | 0.028 | 0.012 | 0.021 | 0.55 | 0.15–0.20 |
| Discovery, missed | 0.19 | 0.020 | 0.010 | 0.013 | 0.44 | 0.16–0.18 |

**Table N5.** Canonical BHB fraction by tRNA region (sequence families, discovery and independent panels combined, ±3-nt window).

| Status | Region | Families | Observed | Host | Position | Dinuc. |
|---|---|---|---|---|---|---|
| called | D-arm | 297 | 0.51 | 0.061 | 0.053 | 0.045 |
| called | anticodon stem | 300 | 0.53 | 0.062 | 0.053 | 0.036 |
| called | anticodon loop | 886 | 0.56 | 0.064 | 0.043 | 0.078 |
| called | variable region | 177 | 0.48 | 0.057 | 0.041 | 0.043 |
| called | T-arm | 456 | 0.77 | 0.090 | 0.041 | 0.121 |
| called | other | 35 | 0.62 | 0.073 | 0.043 | 0.072 |
| missed | D-arm | 437 | 0.33 | 0.053 | 0.045 | 0.051 |
| missed | anticodon stem | 87 | 0.20 | 0.076 | 0.044 | 0.036 |
| missed | anticodon loop | 31 | 0.45 | 0.042 | 0.032 | 0.073 |
| missed | variable region | 128 | 0.21 | 0.062 | 0.052 | 0.037 |
| missed | T-arm | 139 | 0.63 | 0.093 | 0.056 | 0.111 |
| missed | other | 9 | 0.11 | 0.011 | 0.078 | 0.022 |

**Gains.** For census-based gains, observed and null values of member insertions were averaged within gains, gains within loci, and loci were weighted equally (Table N6). For the annotation-based calls, each intron was evaluated at its tRNAscan-SE boundary in the fully spliced context of its host gene; host reassignment was stratified by intron position (canonical 37/38 or noncanonical) and intron-length bin (2,000 permutations), dinucleotide-shuffle and within-gene position nulls (introns moved at least 3 nt, rather than the 8 nt used for census sequences, because tRNAscan-SE boundaries are exact) used 200 replicates, and values were equally weighted over dependency units (Table N7). With the relaxed class included, 95% of the 127 annotation-based calls are BHB-compatible (98% with dependency units weighted equally, against 31% after host reassignment); this permissive metric is shown for comparison only.

**Table N6.** Canonical BHB fraction of census-based gains, equally weighted over loci.

| Gain set | Gains | Loci | Observed | Host | Position | Dinuc. | $P$ |
|---|---|---|---|---|---|---|---|
| all gains (Tier 1) | 129 | 65 | 0.55 | 0.063 | 0.035 | 0.084 | $< 10^{-4}$ |
| all gains (Tier 1 + 2) | 173 | 88 | 0.57 | 0.059 | 0.041 | 0.082 | $< 10^{-4}$ |
| gains with a missed insertion | 23 | 19 | 0.30 | 0.058 | 0.031 | 0.058 | 0.007 |
| gains with only missed insertions | 14 | 11 | 0.26 | 0.044 | 0.028 | 0.027 | 0.064 |

**Table N7.** Canonical and any-class BHB fractions for the annotation-based gain calls, equally weighted over dependency units. Host-reassignment $P = 0.0005$ (the minimum attainable) for every set; dinucleotide and position $P = 0.005$ (minimum with 200 replicates).

| | | | Canonical BHB | | | | Any BHB-like | |
|---|---|---|---|---|---|---|---|---|
| Call set | Calls | Units | Native | Host | Dinuc. | Position | Native | Host |
| all 127 calls | 127 | 8 | 0.59 | 0.080 | 0.032 | 0.005 | 0.98 | 0.31 |
| remaining, exact | 107 | 11 | 0.68 | 0.104 | 0.030 | 0.013 | 0.97 | 0.34 |
| remaining, ±1 nt | 98 | 14 | 0.53 | 0.098 | 0.034 | 0.020 | 0.91 | 0.33 |
| remaining, ±3 nt | 83 | 16 | 0.58 | 0.092 | 0.030 | 0.024 | 0.91 | 0.29 |

#### Supplementary Note 4. Independent panel, sequence reuse, mobile elements and EndA

**Independent panel.** Table N8 lists family means. The selection rule and accessions were fixed before tRNA analysis. The 30 genomes represent 21 GTDB families (2 focal, 19 non-focal); the two Odinarchaeaceae genomes, selected in different strata, form one family. All eight focal genomes are draft assemblies (quality tier GQ-C), which supports presence but not reliable absence. With only two focal families, resampling-based confidence intervals are not informative, so the family-level comparison is reported as an exact permutation probability over all $\binom{21}{2} = 210$ assignments of the focal label.

**Table N8.** Independent panel: intervening sequences per tRNA gene by family (non-pseudogene tRNA genes). Focal families in bold.

| Family | Genomes | Annotated | All (census) |
|---|---|---|---|
| **Freyrarchaeaceae** | 6 | 1.58 | 2.42 |
| **Jordiarchaeaceae** | 2 | 1.50 | 2.01 |
| Borrarchaeaceae | 1 | 0.93 | 1.65 |
| DXJG01 | 1 | 1.32 | 1.60 |
| Odinarchaeaceae | 2 | 1.25 | 1.52 |
| HEL-GB-A | 1 | 0.93 | 1.40 |
| Kariarchaeaceae | 1 | 1.09 | 1.12 |
| Hodarchaeaceae | 1 | 0.79 | 1.11 |
| Wukongarchaeaceae | 1 | 0.77 | 1.10 |
| Hermodarchaeaceae | 2 | 0.77 | 0.98 |
| Thorarchaeaceae | 1 | 0.52 | 0.86 |
| SOKP01 | 1 | 0.45 | 0.86 |
| Sigynarchaeaceae | 1 | 0.55 | 0.86 |
| JABLTI01 | 1 | 0.74 | 0.81 |
| MK-D1 | 1 | 0.41 | 0.78 |
| Baldrarchaeaceae | 1 | 0.61 | 0.74 |
| JAJRWK01 | 1 | 0.62 | 0.64 |
| Heimdallarchaeaceae | 2 | 0.50 | 0.53 |
| Njordarchaeaceae | 1 | 0.48 | 0.50 |
| S146-22 | 1 | 0.32 | 0.42 |
| Panguiarchaeaceae | 1 | 0.36 | 0.42 |

**Sequence reuse.** Within each genome, intervening sequences were joined by single linkage when their alignment identity (matches over aligned length, end overhangs excluded, denominator at least 90% of the longer sequence) reached 70%, 80% or 90% and their lengths differed by $\leq$20%. The statistic is the fraction of sequences in families spanning $\geq$2 tRNA isotypes. The same procedure was applied to dinucleotide-shuffled sequences (two replicates) and to one genome per merged focal lineage (Figure S4). A complementary analysis of 282 published Thermoproteales introns and the annotated introns of the three complete Asgard genomes (complete-linkage clustering at 70% identity) found 26 recurrent families in Thermoproteales and one weak cross-isotype family in Asgard, which disappeared at stricter thresholds.

**Genome quality, mobile elements, reverse transcription and EndA.** Genome-quality tiers combined completeness, contamination, contiguity, N50 and locus recovery; fragmented genomes contributed presence but not absence. Mobile-element context around each locus was scored from hallmark-protein homology (proteins predicted with Prodigal 2.6.3), exact mapping of the 18 mobile elements published for the complete Atabeyarchaeia genomes, local composition, direct repeats and partial-tRNA fragments; geNomad was not rerun, and its published output was used only to define the 18 benchmark elements. Hallmark-protein homology recovered 14 of the 18 elements (15 when the published geNomad calls are added), and a *Caldiarchaeum* tRNA-disruption positive control was recovered. Each of the 71 annotation-based calls at well-assembled loci was compared with three same-genome control tRNA genes matched for isotype, anticodon, local gene density and distance to the contig end; 196 of the 213 controls were intron-free. The donor search compared each gain intron with the 18 published elements (fuzzy partial alignment on both strands) against 100 mononucleotide shuffles per intron. The RNA/cDNA analysis compared encoded-CCA direction in 94 paired loci, searched for local partial-tRNA fragments, and located three credible reverse-transcriptase genes. EndA homologues were identified against $\alpha$ and $\beta$ structural references; 58 of 60 genomes carried an $\alpha$-like protein, 44 two full-coverage paralogues and 16 were unresolved (Figure S6). Across the panel, divergence from the *Aeropyrum* reference was strongly associated with annotated burden ($\rho = -0.75$, adjusted $P = 7 \times 10^{-11}$), largely reflecting shared ancestry; after within-family centring and Benjamini–Hochberg correction no feature was associated with annotated burden (minimum adjusted $P = 0.112$). We repeated the test with census-corrected burden (Supplementary Table S13): across genomes, sequence divergence appeared associated within families ($n = 31$ genomes, adjusted $P = 0.002$), but the signal came from 13 near-identical Freyrarchaeaceae assemblies; after merging near-identical genomes, neither loop length ($\rho = -0.52$, $P = 0.06$) nor divergence ($\rho = -0.44$, $P = 0.11$) was significant (unadjusted; 14 lineages in four families).

#### Supplementary Note 5. Reproducibility

The workflow has two stages. The upstream stage (`scripts/`) downloads the genomes by accession, predicts tRNAs with tRNAscan-SE and ARAGORN, aligns mature tRNAs to the covariance model, matches loci, infers the species tree, makes the annotation-based gain calls and runs the mobile-element, reverse-transcription and EndA screens. The census stage (`analysis/`) starts from the outputs of the upstream stage and regenerates every number in the main text and in this document (`key_numbers.json`), Supplementary Tables S1–S14 and all figures with `bash run_all.sh` in about five minutes on a laptop. Every random procedure uses a recorded seed. Supplementary Table S14 gives SHA-256 checksums of every input and result file and the software versions used for the census stage; environment files for the upstream stage are included in the archive.

### Supplementary Figures

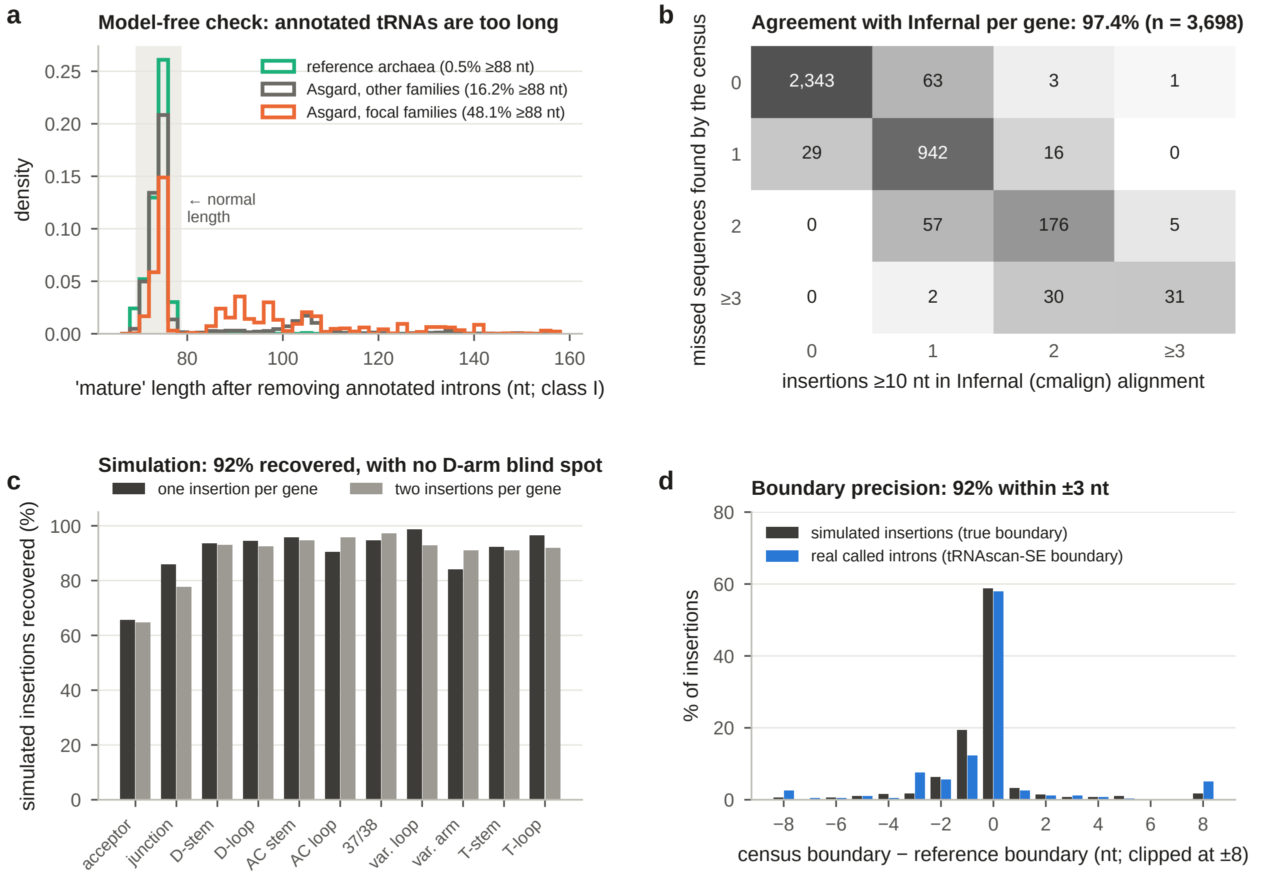


**Figure S1. Independent checks of the census. a**, Model-free check: length of class I tRNAs after removal of the introns annotated by tRNAscan-SE (shaded: normal range); percentages give the share ≥88 nt. **b**, Gene-by-gene comparison with the Infernal covariance-model alignment of "mature" sequences: rows, missed sequences found by the census; columns, insertions of ≥10 nt in insert or local-end states; numbers are genes (discovery and independent panels). **c**, Simulation: recovery of real intervening sequences inserted at random positions of compact tRNA genes, by cloverleaf region, with one or two insertions per gene. **d**, Boundary precision for simulated insertions (true boundary known) and for real annotated introns (relative to the tRNAscan-SE boundary).

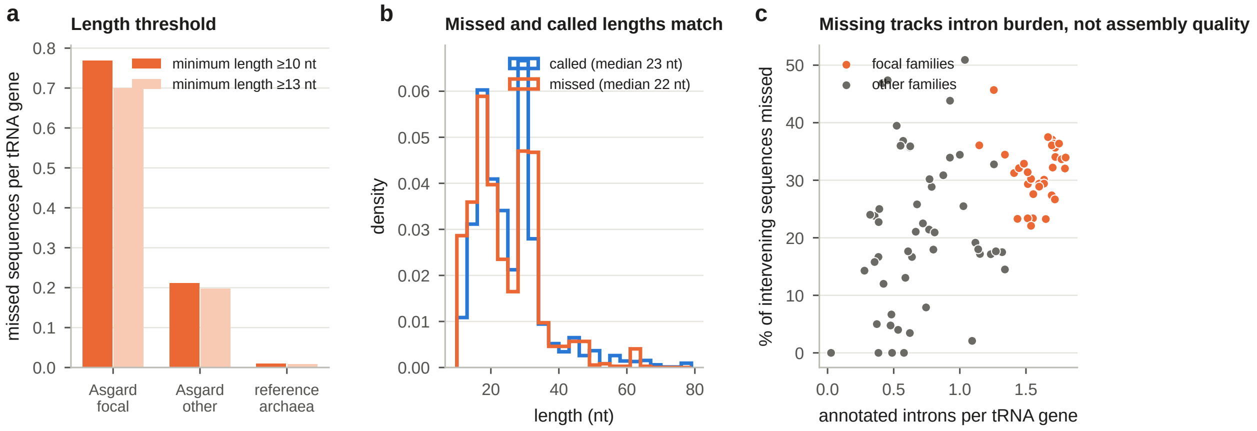


**Figure S2. Robustness of the census.** **a**, Missed intervening sequences per tRNA gene with minimum lengths of 10 and 13 nt. **b**, Length distributions of called and missed intervening sequences (discovery panel). **c**, Percentage of intervening sequences missed by tRNAscan-SE against annotated introns per tRNA gene, for the 90 Asgard genomes (Spearman $\rho = 0.47$, $P = 3 \times 10^{-6}$; against contig number, $\rho = 0.018$; see Supplementary Note 1 for correlations on dereplicated units).

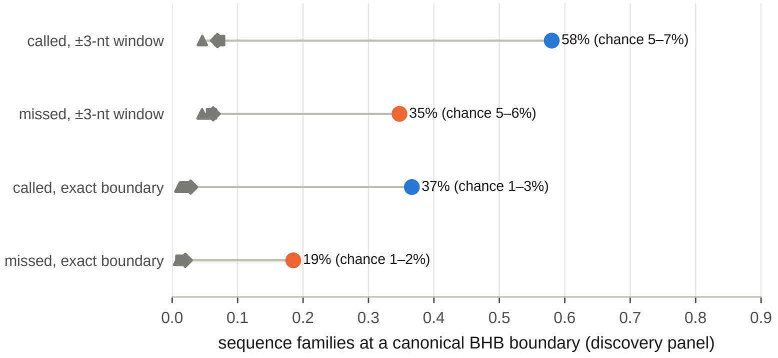


**Figure S3. Placement window.** Canonical BHB fraction for called and missed sequence families (discovery panel) with the ±3-nt placement window and at the exact census boundary; grey symbols show the host-reassignment (diamond), within-gene position (triangle) and dinucleotide-shuffle (square) nulls under the same rule.

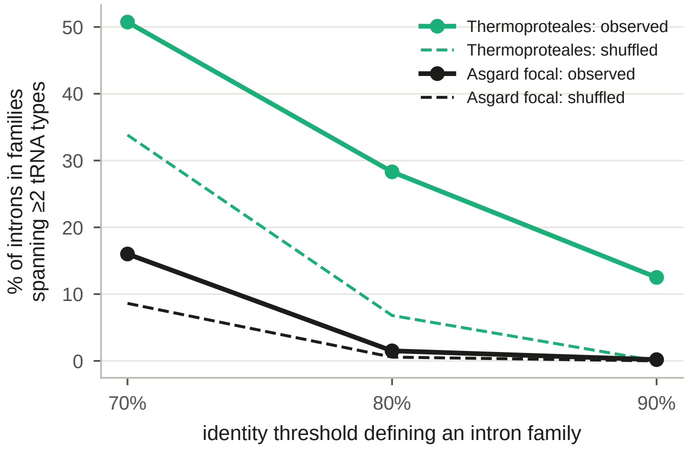


**Figure S4. Sequence reuse across tRNA types at three identity thresholds.** Percentage of intervening sequences in within-genome families that span two or more tRNA isotypes, for Thermoproteales (8 genomes) and focal Asgard lineages (12), observed (solid) and for dinucleotide-shuffled sequences (dashed). At 70% identity chance matches dominate; at 80% and 90% Thermoproteales retain a strong excess and Asgard does not.

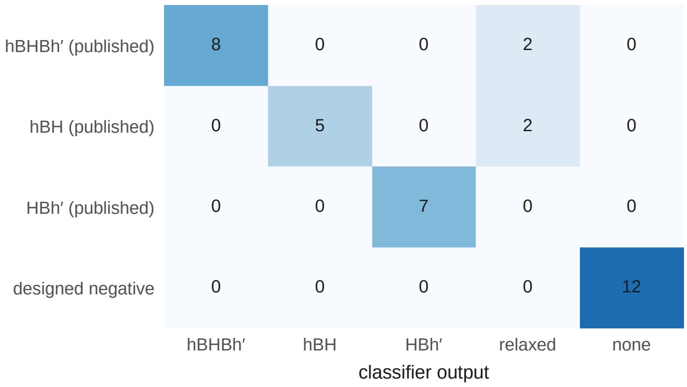


**Figure S5. Benchmark of the BHB classifier.** Published motif classes for 24 archaeal splice junctions and 12 composition-preserving designed negatives (rows) against classifier output (columns). Four published canonical motifs are conservatively assigned to the relaxed class; no negative is called compatible.

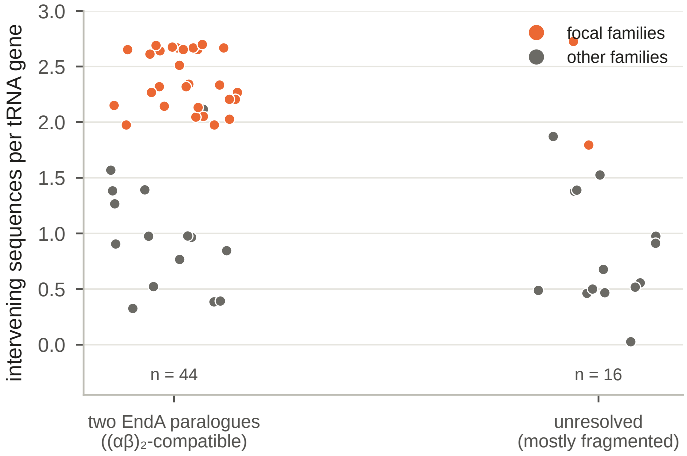


**Figure S6. EndA architecture and intron burden.** Intervening sequences per tRNA gene for discovery genomes with two resolved EndA paralogues compatible with an $(\alpha\beta)_2$ enzyme and for genomes whose EndA inventory could not be resolved. High-burden genomes occur in both categories.

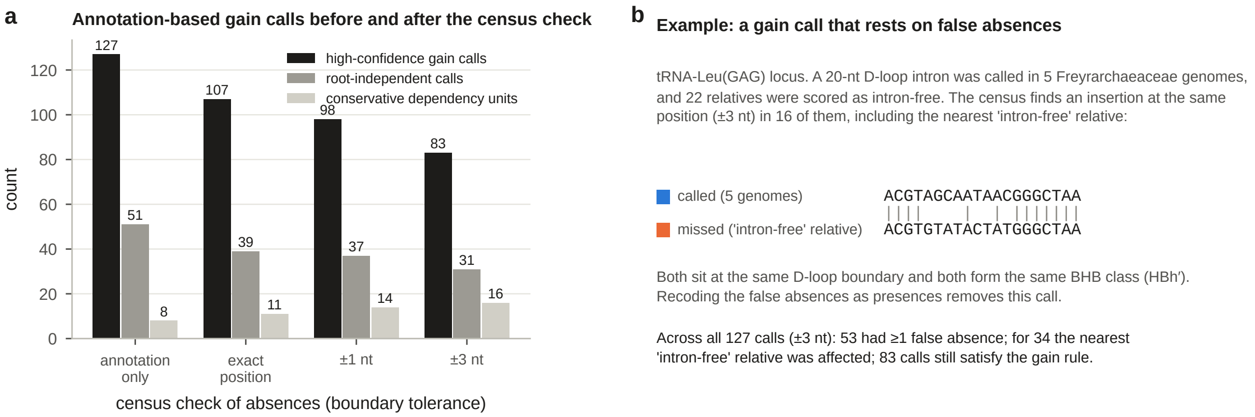


**Figure S7. False absences behind annotation-based gain calls. a**, Annotation-based gain calls, root-independent calls and dependency units, using annotation alone and after recoding census-detected false absences at three boundary tolerances. Units increase as calls are removed because links between otherwise unrelated calls are broken. **b**, An example: the called 20-nt D-loop intron of a tRNA-Leu(GAG) locus (blue) and the missed insertion at the same position in the nearest "intron-free" relative (orange); vertical bars mark identical nucleotides without gaps.

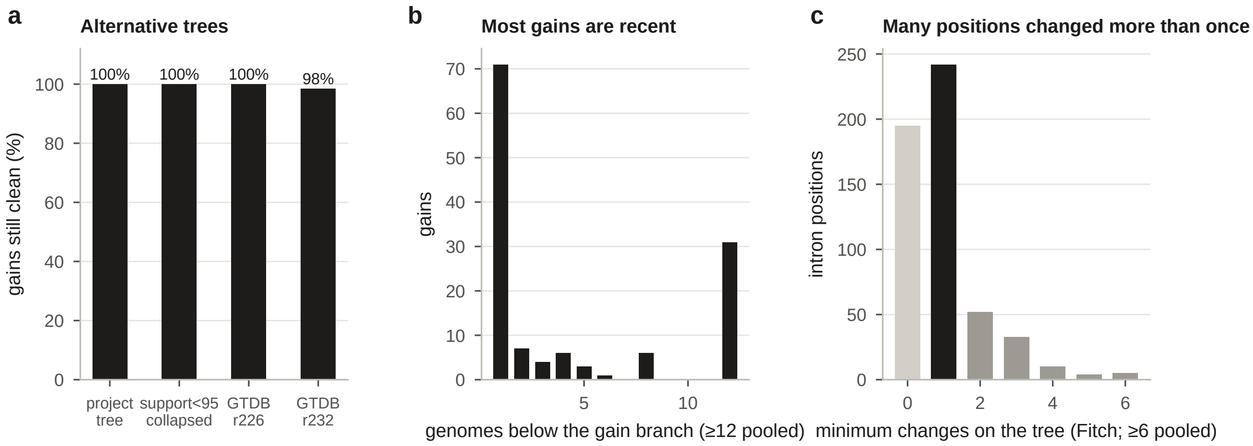


**Figure S8. Census-based gains: robustness and tempo. a**, Percentage of the 129 gains that remain clean (presence confined to one clade with no recovered absence inside it) on the project tree, the project tree with branches of support $<95$ collapsed, and the GTDB release 226 and 232 reference trees. **b**, Number of genomes below each gain branch. **c**, Minimum number of state changes per intron position under Fitch parsimony for all 541 census positions (0, no recovered absence; 1, a single gain or loss explains the pattern; $\geq 2$, repeated changes).

#### Supplementary Tables

Tables are provided as tab-separated files named `Table_S<number>_<content>.tsv`.

| Table | Contents |
|---|---|
| **S1** | Genomes: accessions, taxonomy, assembly quality, panel membership and selection metadata for the 90 Asgard assemblies (GTDB taxonomy) and the 18 reference archaea (NCBI taxonomy). |
| **S2** | tRNAscan-SE intron calls in the discovery and independent panels: panel, locus, alignment column, boundary, canonical status and sequence. |
| **S3** | Census, one row per tRNA gene: annotated, re-found and missed counts, compact reference used, precursor, "mature" and fully spliced sequences. |
| **S4** | Every intervening sequence: status (called or missed), coordinates, length, sequence, cloverleaf region and BHB class (exact boundary and ±3-nt window). |
| **S5** | Independent checks of the census: gene-by-gene comparison with the Infernal alignment, and every simulated insertion. |
| **S6** | Intron burden per genome and per family (`c`, annotated introns; `m`, missed sequences; `n`, tRNA genes; `called_per/allp`, annotated or all intervening sequences per gene), and the independent-panel family test. |
| **S7** | Census character matrix: one row per intron position, with locus, members, annotation status and the state in each of the 60 discovery genomes. |
| **S8** | Census-based gains: rule outcome, branch, root independence, parsimony, state on four trees and BHB values. |
| **S9** | Annotation-based gain calls: rule inputs, topology states, root independence, dependency unit, census check of absences at three tolerances, matching census position and BHB class. |
| **S10** | BHB tests: observed and null values per distinct intervening sequence with its family assignment (window and exact boundary), call-set and gain-set summaries, and tests by tRNA region. |
| **S11** | Sequence reuse across tRNA types, Thermoproteales and focal Asgard lineages, at three identity thresholds. |
| **S12** | Arrival-route tests for each annotation-based gain call (mobile-element tier, RNA/cDNA classification, duplication, sequence mobility). |
| **S13** | EndA inventory, architecture calls and association tests with annotated and census burden. |
| **S14** | SHA-256 checksums of all input and result files, and software versions. |
